# MINI-EX: Integrative inference of single-cell gene regulatory networks in plants

**DOI:** 10.1101/2022.07.01.498402

**Authors:** Camilla Ferrari, Nicolás Manosalva Pérez, Klaas Vandepoele

## Abstract

Multicellular organisms, such as plants, are characterized by highly specialized and tightly regulated cell populations, establishing specific morphological structures and executing distinct functions. Gene regulatory networks (GRNs) describe condition-specific interactions of transcription factor (TF) regulating the expression of target genes, underpinning these specific functions. As efficient and validated methods to identify cell-type specific GRNs from single-cell data in plants are lacking, limiting our understanding of the organization of specific cell-types in both model species and crops, we developed MINI-EX (Motif-Informed Network Inference based on single-cell Expression data), an integrative approach to infer cell-type specific networks in plants. MINI-EX uses single-cell transcriptomic data to define expression-based networks and integrates TF motif information to filter the inferred regulons, resulting in networks with increased accuracy. Next, regulons are assigned to different cell-types, leveraging cell-specific expression, and candidate regulators are prioritized using network centrality measures, functional annotations, and expression specificity. This embedded prioritization strategy offers a unique and efficient means to unravel signaling cascades in specific cell-types controlling a biological process of interest. We demonstrate MINI-EX’s stability towards input data sets with low number of cells and its robustness towards missing data, and we show it infers state-of-the-art networks with a better performance compared to related single-cell network tools. MINI-EX successfully identifies key regulators controlling root development in Arabidopsis and rice, Arabidopsis leaf development, and governing ear development in maize, enhancing our understanding of cell-type specific regulation and unraveling the role of different regulators controlling the development of specific cell-types in plants.

## Introduction

The recent burst in single-cell technologies allowed researchers to study the transcriptional activity within single cells and to describe gene expression profiles characterizing distinct cell-types (Angerer et al., 2017). Despite the initial challenges of the application of single-cell transcriptomics to plant tissues, intrinsic to the nature of their cells (Tripathi and Wilkins, 2021), recent publications have shown their wide applicability (Denyer et al., 2019; Ryu et al., 2019; Jean-Baptiste et al., 2019; Zhang et al., 2019; Shulse et al., 2019; Wendrich et al., 2020; Satterlee et al., 2020; Bezrutczyk et al., 2021; Farmer et al., 2021; Kim et al., 2021; Liu et al., 2021; Zhang et al., 2021a; Zhang et al., 2021b; Marand et al., 2021). For example, Wendrich and colleagues, using a dense single-cell expression atlas of the *Arabidopsis thaliana* root, showed a link between vascular signaling pathways and trichoblast formation during phosphate starvation (Wendrich et al., 2020). Other studies used single-cell transcriptomics to describe detailed developmental trajectories of different cell-types in the Arabidopsis root and vegetative shoot apex (Denyer et al., 2019; Zhang et al., 2021a).

Gene expression information alone, however, might be insufficient to study complex differentiation processes and to characterize how cell-type specific organization is established. One of the main mechanisms contributing to the definition of cell identity involves the coordinated activity of transcription factors (TFs) which, by activation or repression of their target genes (TGs), control gene expression in a tissue-specific manner (Tripathi and Wilkins, 2021). Accordingly, a regulon is defined as the TF and the set of TGs it regulates in a given cell-type. These regulatory interactions are often represented as gene regulatory networks (GRNs), where TFs and TGs are represented as nodes, connected by edges representing their regulatory relationships. As GRNs describe how the expression of defined genes is regulated in specific organs and conditions, they can be mined to identify key molecular players and explore their functions in specific cell-types (Barabási and Oltvai, 2004).

GNRs have been instrumental to elucidate complex plant processes such as response to stress, nutrient signaling, and organ development (Tian et al., 2014; Varala et al., 2018; Chen et al., 2018; De Clercq et al., 2021; Wu et al., 2021). Moreover, these networks have been used to entangle complex regulatory relationships essential for the organization of Arabidopsis’ root tissue, as exemplified by the identification of a series of feed-forward loops regulating secondary cell wall biosynthesis (Taylor-Teeples et al., 2015), and by the identification of broadly expressed master regulators controlling more tissue specific TFs (Brady et al., 2011). The majority of methods used to infer GRNs are purely expression-based, where expression profiles of TFs and TGs are explored and statistically tested to identify regulatory relationships that best explain their expression (Huynh-Thu et al., 2010; Marbach et al., 2012; Matsumoto et al., 2017; Haque et al., 2019; Moerman et al., 2019). While these methods have shed light on regulatory cascades (Ezer et al., 2017; Huang et al., 2018a; Ramírez-González et al., 2018; Denyer et al., 2019), the inferred networks are known to carry many false positives and lack support of physical interactions (Marbach et al., 2012). In order to overcome this problem, several methods explore prior knowledge to identify target genes more accurately, such as chromatin accessibility or transcription factor binding site (TFBS) enrichment (Bonneau et al., 2006; Siahpirani and Roy, 2017; Aibar et al., 2017; Kulkarni et al., 2019; Kamimoto et al., 2020). TFBSs are represented as matches of the specific sequence recognized by the TF (defined as motif) to the underlying DNA sequence.

GRN inference in plants still carries several limitations and challenges, such as the lack of integrated methods combining expression-based networks, characterized by low accuracy, and TF motif information. TF motifs are short and degenerated sequences, thus, the simple mapping of these motifs to genes’ promoter identifies both functional as well as false positive binding sites (Kulkarni and Vandepoele, 2020). To alleviate this problem, enrichment analysis to identify overrepresented TFBSs on a set of co-regulated genes is usually performed (Kulkarni et al., 2018). The relatively low abundance of characterized motifs limits the information available for several TFs, representing another limitation for GRN inference in plants. Finally, although great progress has been made in describing fundamental regulatory relationships between TFs and their target genes (Taylor-Teeples et al., 2015; Varala et al., 2018; Chen et al., 2018; De Clercq et al., 2021), the lack of fine-grained resolution prevents a deeper understanding of cell-type specific regulation. Whereas single-cell profiling represents a promising avenue to address these challenges, validated methods to infer networks from single-cell data in plants are lacking, limiting the characterization of GRNs controlling growth and development of plant organs and cell-types.

To aid GRN inference in plants, we developed MINI-EX, a Motif-Informed Network Inference method based on single-cell EXpression data, which integrates expression-based networks and TF motif information. MINI-EX takes as input a gene-to-cell count matrix and, through a series of sequential steps, extensively tested for parameters optimization using a TF-TG Interactions-Gold Standard, outputs cell-type specific networks and regulons. MINI-EX partially overcomes the challenge of unknown TF motifs by exploiting the similarity of motifs recognized by TFs of the same family. Furthermore, it applies TFBS enrichment to remove putative false positive interactions solely supported by expression data, using binding information obtained from an ensemble approach, which yields increased accuracy for GRN inference (Kulkarni et al., 2019). Finally, different network statistics as well as available gene function information and expression specificity are leveraged to identify and prioritize regulons relevant for a biological process or a cell-type of interest. We demonstrate the applicability of MINI-EX on three recently published root single-cell RNA-Seq datasets of *Arabidopsis thaliana* and *Oryza sativa*, on an Arabidopsis leaf single-cell RNA-Seq dataset, and on a developing ear dataset of *Zea mays* (Denyer et al., 2019; Wendrich et al., 2020; Xu et al., 2021; Kim et al., 2021; Liu et al., 2021). MINI-EX identified regulons for all reported cell-types in Arabidopsis, rice and maize, respectively, and succeeded to accurately prioritize key regulons based on their network characteristics. While we observed many known tissue-specific TFs within the top scoring cell-type specific regulons, several unknown regulators were identified, representing promising candidates for future functional characterization.

## Results

### Inference of cell-type specific gene regulatory networks using MINI-EX

We developed MINI-EX, an integrative method that combines the information obtained from expression profiles deriving from single-cell data and TF motif information to infer cell-type specific GRNs, which can be easily run as a Nextflow pipeline using a Singularity container, offering a portable and reproducible way properly handling dependencies. To evaluate the power and applicability of MINI-EX, we first applied it on a recently published dataset of *A. thaliana* root covering 15,411 cells and 19,241 expressed genes (Wendrich et al., 2020). During the processing of scRNA-Seq data, one of the fundamental steps is the clustering of cells based on similar expression profiles, which are then leveraged to identify cluster-specific genes (upregulated genes) contributing to the definition of cluster identities. We identified 41 single-cell clusters representing 14 cell-types (Wendrich et al., 2020) and generated networks for all the described cell-types.

MINI-EX applies three main steps to infer GRNs starting from single-cell RNA-Seq (scRNA-Seq) data (Figure 1A). First, gene expression profiles are used to infer an initial expression-based network where TFs are associated to putative target genes. Secondly, for each TF, the predicted target genes are tested for enrichment of TFBSs to refine the regulons (TF + TGs) and remove interactions lacking TFBS support. This step adds additional evidence in support of the predicted TF-TG interactions and helps overcoming the problem related to the high false-positive rate of expression-based networks (Marbach et al., 2012). In a third step, gene expression is again exploited in two subsequent phases, where regulons are first filtered based on the TGs’ expression, and later, based on the TF’s expression. More precisely, each regulon is first assigned to a specific cell-type through enrichment analysis based on TGs’ expression, and subsequently these cell-type specific regulons are filtered for the TF’s expression within the cell-type (Figure 1A - step 3a and 3b). This last step allows to define regulons for each cluster of cells and to only retain those controlled by TFs expressed in the same cell-type.

**Figure 1.**
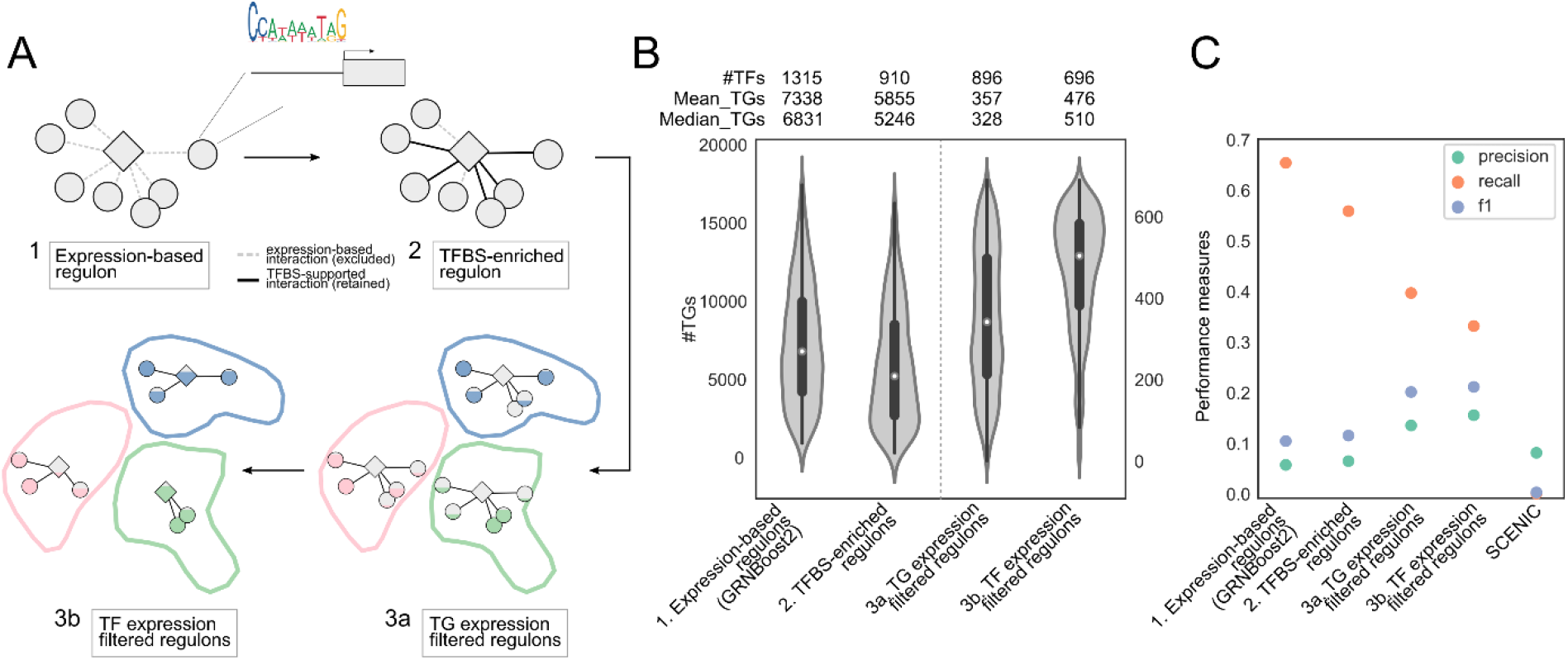
MINI-EX overview and benchmark. (A) Overview of the three steps applied in MINI-EX. Briefly, expression-based regulons composed of TF and putative TGs are computed and successively filtered for TFBS enrichment in the TGs’ regulatory regions, TGs’ and TFs’ expression across single-cell clusters. In the graphical illustration TFs are reported as diamonds and TGs as circles. Step 1 shows an expression-based regulon where interactions are indicated by dashed lines, while in step 2 interactions supported by TFBS enrichment (retained) are indicated by a solid line. The TFBS enrichment is illustrated by the mapping of the TF motif to the gene’s promoter. In step 3a the regulons are filtered based on the TGs’ expression within the single-cell clusters, while in step 3b the regulons are filtered according to the TF’s expression within the single-cell cluster. The expression of TGs and TFs within the specific single-cell cluster is represented by the filling of the circles and diamonds, respectively, while the single-cell clusters are represented by the colored shapes. (B) Violin plots showing the distribution of TGs per each regulon in each of the MINI-EX steps. On the top of each violin plot the number of TFs, the mean and median TGs per regulon is reported. In step 1 and 2 regulons are not yet split in single-cell clusters but a single regulon per TF is reported. (C) Point plot illustrating performance measures (precision, recall and F1 score in green, orange and violet, respectively) for each of the MINI-EX steps and SCENIC.

Although in the past years great effort has been made to determine TF motifs (Jin et al., 2017; Lambert et al., 2019; Fornes et al., 2020; reviewed in Kulkarni et al., 2018), we still lack motif information for nearly half of all Arabidopsis TFs. To avoid discarding initial regulons containing TFs for which motif information is lacking, we allowed enrichment of TFBSs either relative to a specific TF motif or to a TF-family in the second step of MINI-EX. The latter is based on the observation that plant TFs exist in large gene families where often different members recognize highly similar motifs (Weirauch et al., 2014; O’Malley et al., 2016; Wilkins et al., 2016). This allows us to study regulatory relationships for over 80% (1877/2276) of all Arabidopsis TFs.

To ensure optimal prediction performances and establish appropriate cutoffs, each step of MINI-EX was benchmarked against a gold standard obtained by gathering data from ChIP-Seq, Y1H experiments, and manually curated collections, enriched for root-specific genes (82,303 protein-DNA interactions covering 115 TFs, named: “Interactions-Gold Standard”, see Methods) (Supplemental Figure 1). While the TFBS enrichment step allows the refinement of putative target genes resulting in an average reduction of 20% TGs per regulon, the successive steps further reduced TGs by 90% (Figure 1B). This drastic reduction was accompanied by an improved performance represented by a higher F1 score (the harmonic mean of precision and recall), demonstrating that these filtering steps allow us to better control false positives. In fact, while recall dropped by 49% going from the initial general expression-based network to the TFBS-enriched cell-type specific networks, precision and F1 score increased 2.7 and 2.1 fold, respectively (Figure 1C).

We compared the MINI-EX network with a network learned using GRNBoost2, one of the top performing methods for GRN inference solely using single-cell expression data (Moerman et al., 2019; Pratapa et al., 2020), also used in the first step of MINI-EX. We found that the integration of TF binding information greatly improves GRN inference, as 79 out of 91 TFs which passed the TFBSs filtering step showed a higher F1 (Supplemental Figure 2A, green and black lines showing increase in F1 score after TFBS enrichment). Globally, the networks inferred by MINI-EX showed a higher precision compared to GRNBoost2 (15% and 5.6% for MINI-EX and GRNBoost2, respectively), and two times higher F1 (21% and 10% for MINI-EX and GRNBoost2, respectively) (Figure 1C). Alongside, we compared the performances of MINI-EX with SCENIC, another single-cell network inference method (Aibar et al., 2017). Similarly to MINI-EX, SCENIC uses scRNA-Seq data and TF motif information to infer regulons. As SCENIC only supports data analysis for human, mouse and fly, we constructed an Arabidopsis motif mapping file (see Methods) to evaluate its performance. We observed that MINI-EX globally performed superior to SCENIC (Figure 1C, Supplemental Figure 1). While the initial expression-based networks inferred by both methods showed similar F1 scores (10.3% MINI-EX, 9.7% SCENIC, Supplemental Figure 2), the performance of SCENIC on the final TFBS-enriched regulons dropped below 0.5% (Figure 1, Supplemental Figure 1). Moreover, MINI-EX identified 63 more regulons from the Interactions-Gold Standard compared to SCENIC, and showed a higher F1 score for the majority of the regulons inferred by both methods (12/18 regulons showed a higher F1 in the MINI-EX inferred networks, Supplemental Figure 2B).

The dataset we used for testing MINI-EX contains >15,000 cells, a higher number compared to other published studies which profiled between 3000 and 7500 cells (Denyer et al., 2019; Ryu et al., 2019; Jean-Baptiste et al., 2019; Zhang et al., 2019). To validate if MINI-EX performs equally well for smaller datasets, we progressively downsized the initial matrix by randomly selecting sub-samples of the total number of cells (from 100% to 10%, in 10 steps, see Methods). MINI-EX showed a high stability on the down-sampled datasets, as we were able to retrieve 92% (average of 641/696 TFs) of the TFs, and 82% (average of 3556/4340) of the regulons starting from a reduced matrix of ~1500 cells (Supplemental Figure 3A). Additionally, we performed a MINI-EX run on another scRNA-seq root dataset (Denyer et al., 2019) containing 4509 cells, yielding 2552 regulons for a total of 637 TFs. Despite the reduced size of the dataset and the inferred network, 88% (561/637) of the TFs were common to the larger dataset (Wendrich et al., 2020), and performance measures were largely similar (precision: 15.4% and 17.2%, recall: 33% and 25.2%, and F1: 21% and 20.5% for Wendrich and Denyer datasets, respectively; Supplemental Figure 3B), indicating that MINI-EX also obtains good performance for smaller datasets.

An intrinsic property of scRNA-Seq is the sparsity of the data related to the presence of many missing values (dropouts) within the gene-to-cell expression matrix attributable to either technical noise or true absence of signal (Hicks et al., 2018). To mitigate the negative effects missing values have on downstream analyses, several imputation methods have been developed. To assess whether the imputation of missing values also improves GRN inference performance, we tested two top-performing methods, MAGIC and SAVER (Huang et al., 2018b; van Dijk et al., 2018; Hou et al., 2020). Both methods showed a lower F1 score compared to the networks inferred using the non-imputed matrix, albeit SAVER showed a slightly higher precision (16%, 14.3%, 15.4% for SAVER, MAGIC and MINI-EX, respectively), indicating that it recovers a higher fraction of true positives over the total number of predictions (Supplemental Figure 3C). Overall, these results reveal that MINI-EX is a robust method to learn GRNs starting from available single-cell RNA-Seq datasets.

MINI-EX inferred networks for all the identified single-cell clusters except *unknown, putative Lateral Root Cap (LRC)-0*, and the distribution of the 696 TFs across the 40 single-cell clusters was visualized as a clustermap reporting the number of TGs controlled by each TF (Figure 2A). A total of 4340 regulons were inferred (Supplemental Dataset 2) describing the regulation of 8772 TGs. The dendrogram for the single-cell root clusters showed three main groups: one composed by the LRC clusters, another including meristematic clusters such as initials, dividing, Quiescent Center (QC), and cortex, and the final one comprising clusters related to stele (procambium, pericycle and phloem) and endodermis. While the dendrogram highlights how clusters of related cell-types tend to share similar regulatory programs, we observed only one of the three clusters representing endodermis (29) grouping with the stele clusters, suggesting a regulatory differentiation of these endodermis clusters. MINI-EX leverages cluster-specific expression to assign regulons to different cell-types. A detailed analysis of the composition of these cell clusters showed that although the number of upregulated genes per single-cell cluster is highly variable (minimum of 23 in *pericycle-16.0* to a maximum of 1418 in *initials-10*, Figure 2B), all clusters showed more than 50% of their upregulated genes to be part of regulons, either as TGs or TFs, with 8/41 clusters for which all upregulated genes were part of regulons (Figure 2B, genes part of regulons as TGs and TFs in orange and red, respectively). Globally, 5.6% of the upregulated genes were assigned as the main regulator (TF) of at least one regulon, while 89% was represented as TG.

**Figure 2.**
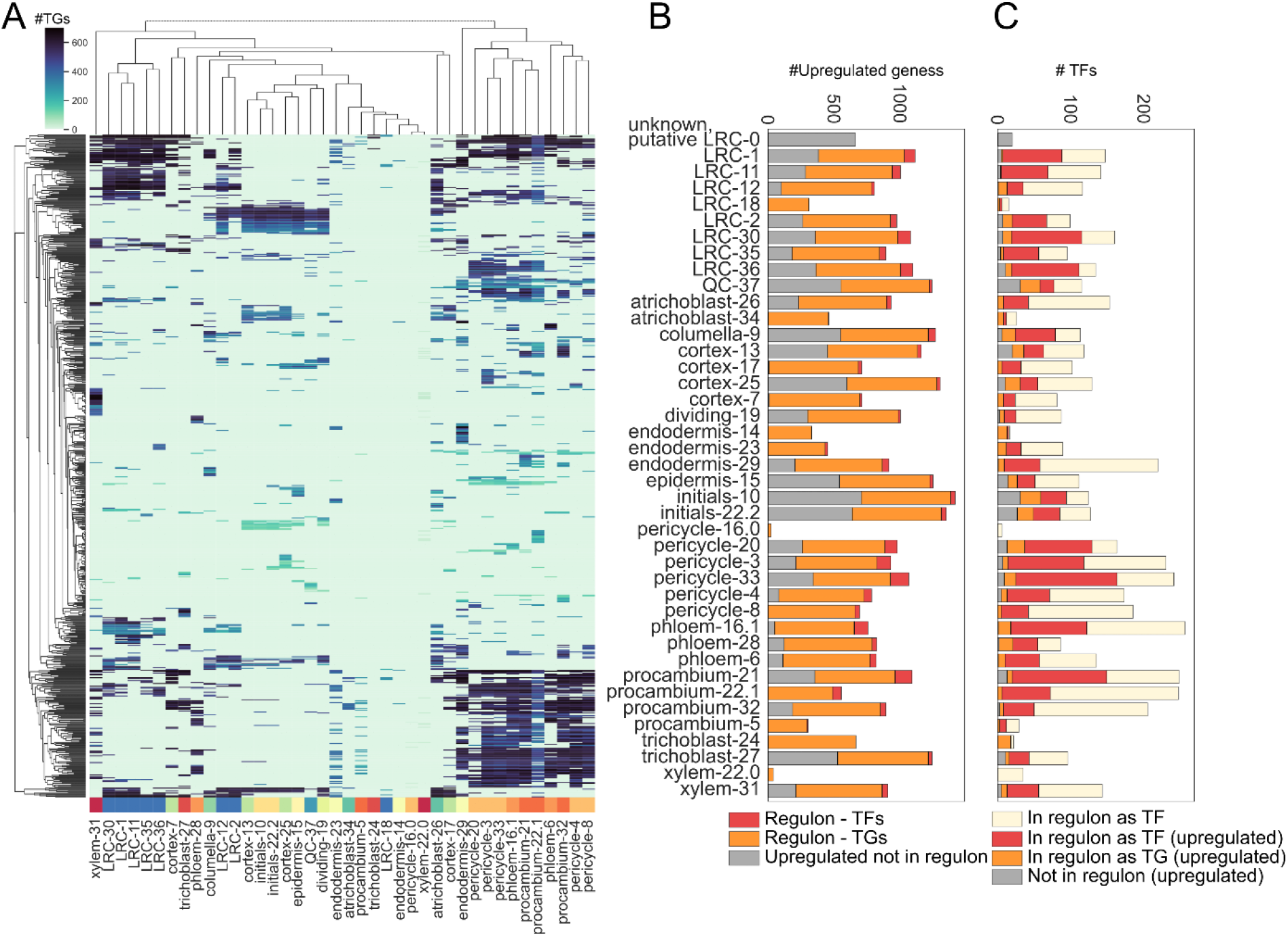
Regulon distribution and composition across the different cell clusters. (A) Clustermap displaying the distribution of 4340 regulons (y-axis) across the identified single-cell clusters (x-axis). The color of the cells represents the number of TGs of each regulon. The different single-cell clusters are color-coded according to their identity. (B) Stacked bars reporting the number of upregulated genes for each single-cell cluster. Red and orange indicate upregulated genes which are part of regulons as main regulator or TG, respectively. (C) Stacked bars showing the number of TFs part of regulons (yellow, red and orange) or just upregulated (gray) for each single-cell cluster.

We preferred not to limit MINI-EX to only return regulons for which the regulator is itself upregulated within the enriched single-cell cluster, as we reasoned that for a TF to regulate its TGs it might be sufficient to be expressed in a substantial fraction of cells within the cluster (Supplemental Figure 1). This resulted in the identification of regulons for clusters that did not show any TFs among their upregulated genes, but are yet, relevant for the cell-type. Examples of these are several VNDs, known to control xylem differentiation (Kubo et al., 2005), in *xylem-22.0*, and NUC in *pericycle-16.0*, known for its involvement in the regulation of vascular pattern divisions (Cui et al., 2011)(Figure 2C, light yellow for TFs controlling regulons and expressed by at least 10% of the cells within the single-cell cluster; Supplemental Dataset 2). Interestingly, cell-types with a meristematic-like function such as initials, QC and columella, showed the largest fraction of TFs not controlling any regulon (Figure 2C, gray). Conversely, several stele clusters (pericycle, procambium and phloem) are among the ones with the highest number of TF regulators (Figure 2C, red). This suggests that, given their role in the Arabidopsis’ root system and their high level of specialization, these stele clusters are under a more complex and coordinated regulation.

In contrast to the original study where only a subset of high-quality cells was retained for downstream analyses, we here included all sequenced cells which passed standard quality filters (see Methods), simulating a standard scRNA-Seq output. A parallel MINI-EX run on the original dataset was performed and showed that, despite the more relaxed filter used on the reprocessed dataset, for 10/13 cell-types > 75% of the regulators inferred from the reprocessed data overlapped with regulators of the same cell-types from the original dataset (Supplemental Figure 4B). A closer inspection of the top 50 regulons per cluster showed how cortex, the cell-type which showed the highest variability, now shows a much larger overlap (Supplemental Figure 4C). Overall, good agreement was observed between clusters of the same identity, with some exceptions where the top 50 TFs showed high overlap with regulators from neighboring tissues, such as *LRC-12* and epidermis, and *LRC-2* and columella. It is worth noticing that low Jaccard Index values characterizing some of the re-analyzed clusters are due to the low number of inferred regulons (<50) (Supplemental Dataset 1).

Taken together, these results demonstrate how MINI-EX, leveraging cluster-specific expression, successfully learns cell-type-specific GRNs which can be further studied to gain insights into regulatory mechanisms governing cell-type development and functions.

### Development of an accurate integrative network-based method to prioritize regulons important for the control of cell-type specific regulation

To evaluate whether MINI-EX successfully retrieves regulators known for their involvement in the control of root development, we compared the TFs of the inferred regulons to a second gold standard obtained by collecting genes with an experimental validated and/or literature supported annotation related to root and vasculature (845 root-related genes including 169 TFs, named: “Functional-Gold Standard”, see Methods). The majority (706) of root-related genes was expressed in our scRNA-Seq dataset and among these, 139 are TFs and 95 (68%) are controlling at least one of the inferred regulons (Figure 3A - “root-related TFs”). Moreover, by including TFs only present as TGs, we could identify 19 additional root-related TFs, resulting in the retrieval of 82% of the TFs part of the Functional-Gold Standard (Figure 3A).

**Figure 3.**
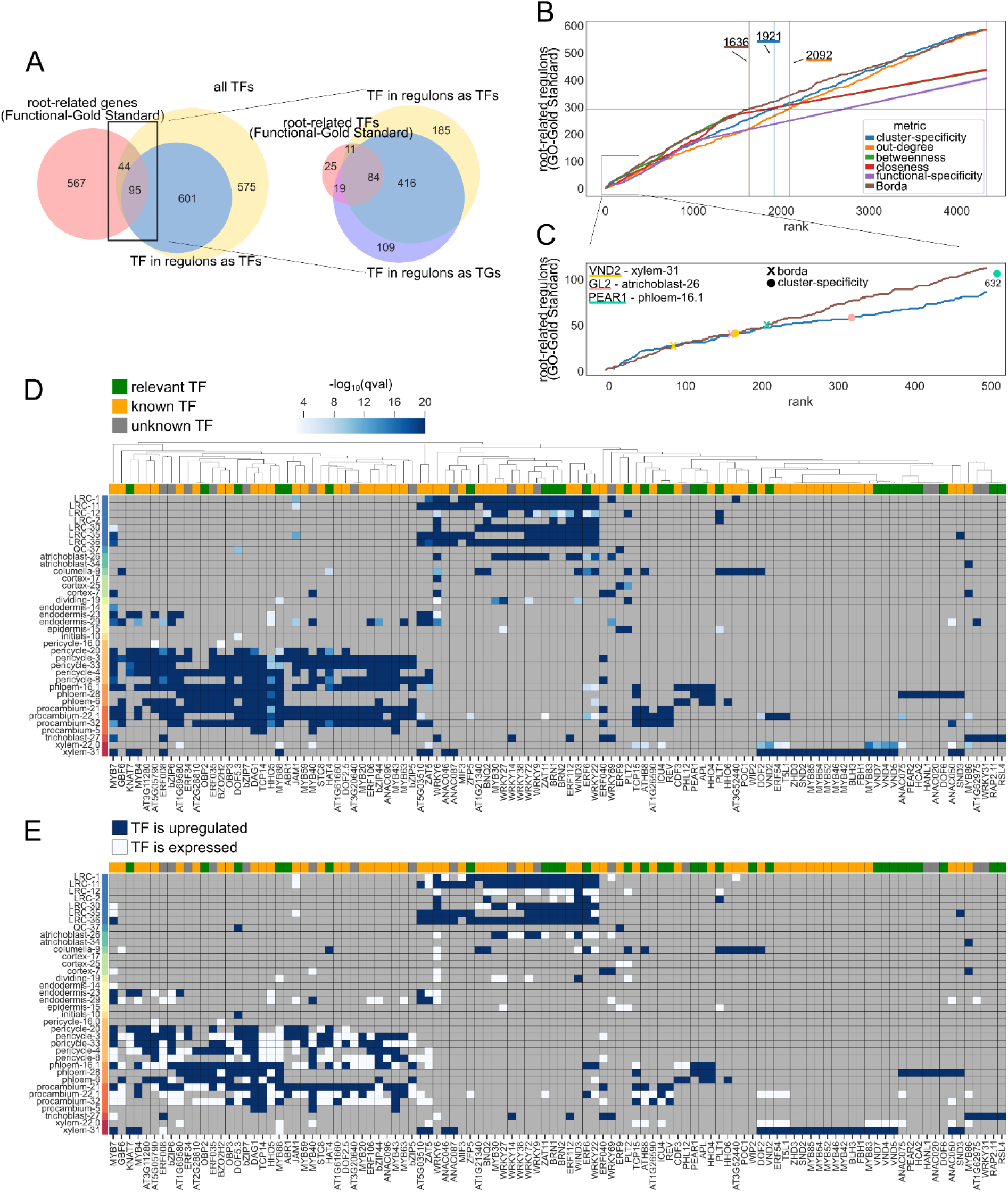
Prioritization of regulons controlling Arabidopsis root architecture. (A) Venn-diagram reporting the overlap of root-related genes from the Functional-Gold Standard with the inferred regulons. (B) Evaluation of prioritization methods. All inferred regulons are ranked based on five single metrics and Borda counts (x-axis). The identification of a root-related regulon (“relevant TFs”, from the Functional-Gold Standard) is shown by the counts on the y-axis. The horizontal line indicates half of the root-related regulons (“relevant TFs”, from the Functional-Gold Standard), while the vertical lines indicate, for each metric, the rank at which 50% of the relevant root-related regulons is retrieved (R50). The six metrics are displayed in different colors as reported by the legend. (C) Enlargement of the first 500 ranks for cluster-specificity and Borda counts in blue and brown, respectively. Three examples of relevant root-related regulons are reported. “X” indicates the ranks given by Borda counts, while “O” indicates the cluster-specificity ranks. (D) Clustermap showing cluster-specificity (the −log_10_(qval) of the single-cell cluster enrichment) of the top 150 inferred regulons. Green, yellow and gray boxes indicate whether the TF controlling the regulon has a root-related GO annotation (“relevant TF”, from Functional-Gold Standard), another GO annotation (“known TF”), or no functional annotation (“unknown TF”), respectively. The different single-cell clusters are color-coded according to their identity. (E) Clustermap showing TF expression related to the top 150 inferred regulons. Blue boxes indicate that the TF is upregulated in the cell cluster is acting, while white boxes indicates that the TF is expressed by at least 10% of the cells belonging to the single-cell cluster. The different single-cell clusters are color-coded according to their identity.

Given the large number of regulons present in the inferred single-cell GRNs, we reasoned that some TFs are more important for controlling root organization compared to others. Therefore, we leveraged the information from network centrality (out-degree, betweenness, closeness) and expression specificity (FDR corrected p-value of cluster enrichment, hereafter referred to as “cluster-specificity”) to prioritize regulons (see Methods). By default, MINI-EX first ranks the inferred regulons based on these four single metrics, and successively uses the geometric mean of these ranks, referred to as Borda counts, to obtain a final ranking. To evaluate this prioritization strategy, we tested at which rank half of the regulons with existing root Gene Ontology (GO) annotations, based on the Functional-Gold Standard, were recovered (called R50 value). The ranking based on Borda counts reached R50 238 ranks earlier than the best performing single metric (1683 and 1921 for Borda and cluster-specificity, respectively, Supplemental Figure 5A) resulting in a better prioritization (12% improvement) of regulons associated with relevant root-related TFs. If a list of expected GO terms is provided by the user, a fifth metric based on the functional enrichment of these GO terms for the regulons’ TGs (hereafter referred to as “functional-specificity”) is introduced, and a customized ranking using weighted Borda counts is performed (see Methods). MINI-EX progressively tests all possible combinations of the five metrics and defines the best prioritization strategy as the one which retrieves half of the relevant regulons fastest (smallest R50). Moreover, MINI-EX provides both a global ranking of the inferred regulons and a cluster-specific ranking, the latter allowing to select important regulons for cell-type clusters of interest. Additionally, an evaluation of the PageRank algorithm revealed that prioritization using cluster-specific Borda ranking outperformed cluster-specific PageRank results (Supplemental Figure 5B).

For the Arabidopsis single-cell root dataset we used, the weighted Borda ranks calculated combining cluster-specificity, functional-specificity, closeness and betweenness centrality, turned out to be the best combination, reaching R50 285 and 456 ranks earlier than the two single best performing metrics (1636, 1921 and 2092 for Borda, cluster-specificity and out-degree centrality, Figure 3B in brown, blue and orange respectively, Supplemental Figure 5C). This corresponds to a 8% and 22% improvement, respectively. To gain more insights into these results, we inspected some of the early-ranking regulons and compared their Borda and cluster-specificity ranks (Figure 3C). For instance, the ranking based on Borda placed the known xylem-specification regulator VND2 (Yamaguchi et al., 2010) in *xylem-31* earlier than the ranking based on cluster-specificity (Figure 3C yellow, ranks 90 and 169 respectively). The same trend was observed for the regulon of the atrichoblast differentiation marker GL2 in *atrichoblast-26* (Figure 3C, pink, ranks 169 and 323 respectively) (Masucci et al., 1996), while a larger difference was observed for PEAR1, known to promote radial growth (Miyashima et al., 2019), in *phloem-16.1* (Figure 3C, turquoise, ranks 212 and 632 respectively).

Taken together, these results demonstrate the power of the prioritization system of MINI-EX, where different combination of metrics based on network properties, expression specificity, and functional enrichment of regulons’ TGs are automatically tested to ensure an optimal prioritization of the inferred regulons towards a tissue or biological process of interest.

### Identification of known and novel tissue-specific TFs among the prioritized regulators

The integrative prioritization strategy implemented by MINI-EX highlighted, for the analyzed single-cell dataset, relevant regulators known for their role in root development. To evaluate whether MINI-EX also succeeds in identifying novel root-related genes, we searched the database of genome-wide association studies for Arabidopsis (AraGWAS) (Togninalli et al., 2018), for genes which showed a significant association with root phenotypes (named “root-GWAS”). We collected a total of 560 genes, of which 377 (67%) were expressed in our single-cell root dataset. Among these, 25 are TFs and 15 of these appeared as main regulators of the inferred regulons (Supplemental Table 1). The distribution of root-GWAS genes in the regulons’ TGs was assessed and we observed that on average 1.8% of the regulons’ TGs overlapped with the upregulated root-GWAS (Supplemental Table 1). As MINI-EX refines the regulons’ TGs based on the upregulated genes within the single-cell cluster, we only focused on the 187 root-GWAS genes which were upregulated in our dataset, and we observed that the majority of these (155) was represented in the regulons’ TGs, and only 48/4340 regulons did not show any root-GWAS gene among their TGs. These results give additional proof that MINI-EX successfully identifies relevant novel candidates which can be further explored for functional characterization studies.

The top 150 prioritized regulons, corresponding to 108 TFs, were further analyzed for their role in root development. Whereas 38 (25.3%) regulons are controlled by TFs which have been experimentally characterized to be involved in root/vasculature organization (part of the Functional-Gold Standard), 85 (56.7%) regulons are controlled by TFs which have an experimentally validated functional annotation unrelated to root biology, and 27 (18%) do not have any experimentally validated function (Figure 3D, 3E, green, yellow and gray squares named “relevant TF”, “known TF”, and “unknown TF”, respectively). Moreover, four TFs of the top 150 regulons, bZIP6, BZO2H2/bZIP9, KNAT7 and HAT4, showed to have a root-GWAS association. We visualized the top-scoring regulons as a clustermap showing their expression specificity (cluster-specificity) across the different cell-types (Figure 3D). This metric demonstrated to be of great importance in the regulon prioritization process (Supplemental Figure 5A), and cell-type specific expression for the majority of regulons was indeed observed. Among them, ten regulons well known for their involvement in phloem development (green, Supplemental Figure 6A), such as APL, OBP2, PEAR1 and PEAR2 (Bonke et al., 2003; Skirycz et al., 2006; Miyashima et al., 2019), were controlling *phloem-28* and *phloem-6*. Similarly, five known xylem regulons, comprising VND2, 4, 5 and 7 were identified as regulators for xylem (*xylem-31* and *xylem-22.0* - green, Supplemental Figure 6A) (Kubo et al., 2005). Other interesting examples included BRN1 and BRN2, known for their role in lateral root cap maturation (Bennett et al., 2010), which were found to control several LRC clusters, and RSL4, responsible for root hair cell growth (Datta et al., 2015), found to control *trichoblast-27*.

While all the TFs controlling the 150 top ranked regulons showed to be upregulated in the cell-type the regulon was enriched for (Figure 3E, dark blue), we also found examples where the correctly recovered TF was only expressed (in at least 10% of the cells) in the single-cell cluster. For instance, we identified ATHB8, a marker gene for procambial cells (Baima et al., 2001) regulating *procambium-32* and *22.1* (Supplemental Dataset 2). Similarly, genes showing good rankings were BRN1 in *LRC-2* (Bennett et al., 2010) and MYB88, known to be expressed in developing xylem cells (Lei et al., 2015), in *xylem-31* (Supplemental Dataset 2). This confirms that important regulatory interactions are not necessarily characterized by TFs showing differential expression (De Clercq et al., 2021).

Finally, we could evaluate the accuracy of the predicted TGs for 14 TFs of the top 150 regulons by comparing them to the experimentally validated interactions summarized in our Interactions-Gold Standard (Supplemental Figure 6B). Regulators with a good recovery included VND7, which showed an enrichment for known interactions in both xylem clusters, with a recall of 87% for the regulon controlling *xylem-31*. Similarly, MYB63 and MYB20, known to be involved in lignin biosynthesis (Zhou et al., 2009; Geng et al., 2020), showed a significant enrichment and a recall of up to 80% in pericycle and procambium clusters, both related to xylem development (Supplemental Figure 6B, hypergeometric test, FDR corrected p-value < 0.05).

To evaluate the performance of MINI-EX’s prioritization strategy for another organ, we analyzed an Arabidopsis leaf single-cell dataset (Kim et al., 2021) composed of 5938 cells and 10 cell-types. Compared to the root dataset, a higher fraction of the top 150 regulons (52, 34.7%) was represented by regulons controlled by TFs reported to play a role in leaf development (e.g. Zinc finger DOF, MYB, AP2/ERF and TCP TF families), particularly in companion cells (8/16), epidermis (12/21) and guard cells (8/18) (“relevant TF”, green, Supplemental Figure 7), while 86 regulons (57.3%) showed to have a known function unrelated to leaf biology (“known TF”, yellow, Supplemental Figure 7), and 12 regulons (8%) with an unknown function (“unknown TF”, gray, Supplemental Figure 7).

Overall, these results demonstrate how the regulon prioritization successfully identifies relevant cell-type regulators as well as bona fide target genes, but also uncharacterized TFs linked to root biology through GWAS analysis. As a result, the inferred single-cell networks enhance the selection of novel candidates playing a role in controlling root cell-type development in Arabidopsis.

### Single-cell network inference allows to map experimentally validated target genes of three NAC TFs across different root cell-types in rice

As single-cell data is becoming available for other plant species apart from Arabidopsis, we provide MINI-EX support for two other species, *Oryza sativa* and *Zea mays*. To illustrate MINI-EX’s applicability to infer cell-type specific networks in cereals, we analyzed a recently published single-cell dataset of rice root (Liu et al., 2021) composed of 18,927 expressed genes and 13,450 cells, representing 19 clusters corresponding to 8 cell-types. MINI-EX inferred 681 regulons controlled by 220 TFs across all identified clusters, except *cortex-9*. A relatively high cell-type specific regulation was observed, with 85/220 TFs regulating only one cell-type, and the majority was identified in cortex, endodermis and trichoblast (Supplemental Figure 8A). Among the trichoblast-specific regulons OsRHL1, known to be regulating root-hair development in rice (Ding et al., 2009), was identified, together with LOC_Os10g38820, an ortholog of the Arabidopsis SPI gene, also involved in root-hair development (Chin et al., 2021). The network inferred by MINI-EX covered interactions for 220 TFs, representing 23% (220/937) of the total TFs expressed in the rice root atlas. Although reduced TF motif information was available for rice in comparison to Arabidopsis (1244 and 1699 motifs covering 1643 and 1877 TFs, for rice and Arabidopsis, respectively), the TFBS enrichment step reduced the number of studied TFs by 13% (815/937) for rice, against 31% (910/1315) in Arabidopsis (Supplemental Figure 8B). The last step of MINI-EX, where cell-type specific expression is integrated, showed a stronger effect on the reduction of analyzed TFs for rice, where only 27% (220/815) of the previously identified TFs were maintained. This is in contrast with Arabidopsis, where 76% (696/910) of the TFs which passed the TFBS enrichment step were retained in the final network (Supplemental Figure 8B), revealing that the TF expression specificity in the rice dataset is globally lower, resulting in a smaller single-cell network for rice.

To assess the quality of regulatory interactions of the rice root networks, the regulons associated to three NAC TFs, OsNAC5, OsNAC6 and OsNAC9, were explored. The NAC (NAM/ATAF/CUC) transcription factor family is one of the largest family in plants, comprising more than 100 members in Arabidopsis and rice known to be involved in development and stress responses (Shao et al., 2015). In rice, NAC TFs are well-studied for their involvement in stress signaling (reviewed in Crow et al., 2016), but crucial is also their role in development (Jeong et al., 2013; Mathew et al., 2016; Mao et al., 2017; Mao et al., 2020). Among the 83 NAC TFs expressed in the single-cell rice root dataset, 25 were inferred as regulators in at least one single-cell cluster, and their presence in almost all of the identified cell-types suggests a regulatory role in the development of different cell identities (Supplemental Dataset 4). To validate the NAC regulons, experimental data from bulk ChIP- and RNA-Seq performed on three NAC overexpression lines (OsNAC5, OsNAC6 and OsNAC9) of rice roots was collected (Chung et al., 2018). Experimentally validated TGs were defined as genes which were both differentially expressed and bound in the RNA-Seq and ChIP-Seq bulk experiments, respectively, resulting in 125, 222 and 465 TGs for OsNAC5, 6 and 9, respectively. Considering all NAC-associated regulons, we were able to retrieve 15, 35 and 85 experimentally confirmed TGs, corresponding with a rather low recall (between 12 and 18%). MINI-EX leverages the cell-type specific upregulated genes to assign expression specificity of a regulon, and consequently, a TG needs to be upregulated within the specific cell-type to be part of a regulon. When the experimentally validated TGs were filtered for being upregulated in at least one single-cell cluster, relatively to other cell clusters, recall increased to 52, 66 and 78%, respectively (Figure 4A, in parenthesis the experimentally validated genes upregulated in at least one single-cell cluster). When exploring the cell-types regulated by the three NACs, a large ubiquity of expression was observed, especially for OsNAC9, whose regulons showed a significant overlap with its experimentally validated TGs for all single-cell clusters, while conversely, OsNAC6 and 5 only showed a significant overlap for some of the inferred regulons (Figure 4B, “X” indicating a regulon with no significant overlap with the TF’s experimentally validated TGs).

**Figure 4.**
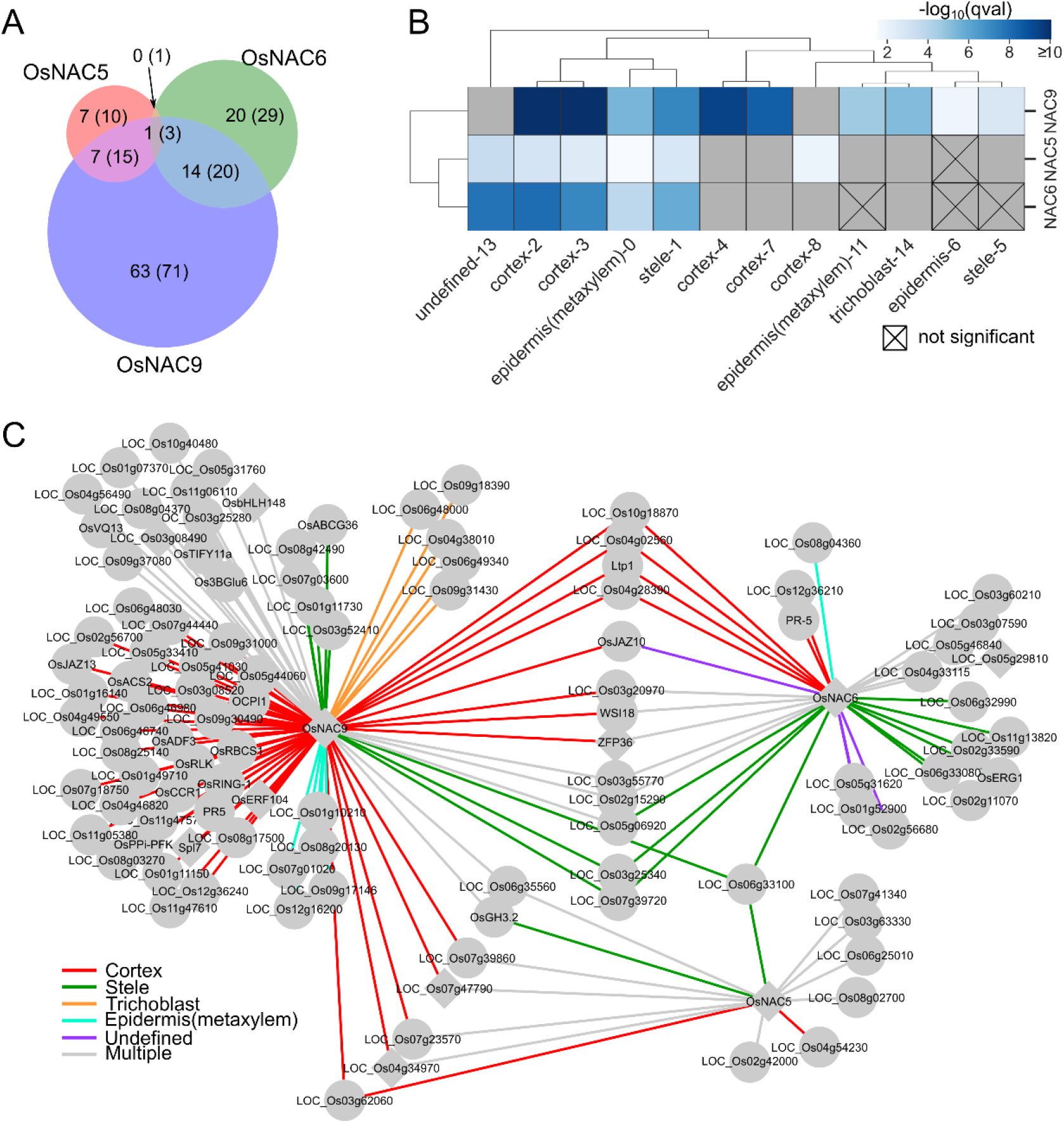
Validation of the inferred cell-type specific NACs regulons. (A) Venn diagram of the overlap between experimentally validated target genes for OsNAC5, OsNAC6 and OsNAC9 regulons. Numbers in parenthesis represent the number of experimentally validated TGs which are upregulated in at least one of the single-cell clusters, numbers outside parenthesis represent experimentally validated TGs of the inferred regulons. (B) Clustermap showing the significance of the overlap (hypergeometric test, FDR corrected p-value < 0.05) between experimentally validated TGs of the three NAC TFs and the predicted TGs in the different single-cell clusters. Gray boxes indicate single-cell clusters for which no OsNAC5, OsNAC6 or OsNAC9 regulon have been inferred (C) GRN of OsNAC5, 6 and 9. Diamonds represent TFs, while circles represent TGs. The color of the edges represents the cell-type the interaction occurs, gray color indicates that the interaction is present in multiple cell-types. Only experimentally validated interactions are reported.

Information from bulk experiments can be coupled with single-cell transcriptomic data to obtain a fine-grained view of specific cell-type regulation which would be otherwise indiscernible. To this end, the cell-type specific networks of the three NACs were further investigated. For the ease of illustration, only the cell-types which showed a significant overlap between experimental and predicted TGs, are depicted (Figure 4B, hypergeometric test, FDR corrected p-value < 0.05). For the TGs present in this network, the majority (72/108) showed to be specifically regulated in a single cell-type, with only 36 confirmed TGs which appeared in regulons of multiple cell-types (Figure 4C, gray edges for interactions occurring in multiple cell-types), indicating a strong cell-type specific regulation. The only gene which showed to be regulated by all three NACs is LOC_Os06g33100 in stele, a peroxidase whose ortholog in Arabidopsis (PRX36) is known to be involved in cell wall degradation during seed development (Francoz et al., 2019). Finally, the large number of regulatory interactions observed in cortex and stele (Figure 4C, red and green edges, respectively) strongly support the reported phenotype, where NAC overexpression lines are characterized by enlarged cortical (OsNAC5) and xylem cells (OsNAC5, 6 and 9) (Jeong et al., 2013; Chung et al., 2018). Taken together, these results show how bulk ChIP-Seq data can be leveraged, together with MINI-EX’s GRN inference on single-cell data, to map cell-type specific regulatory networks and uncover the role of individual regulators.

### Cell-type specific networks recover relevant regulators for maize ear development

GRN inference in maize is known to be challenging because of its large genome size and reduced TF motif information (Zhou et al., 2020; Tu et al., 2020). To evaluate the ability of MINI-EX to infer single-cell GRNs in maize we analyzed a recently published developing ear dataset containing 13,196 cells and reporting 21,704 expressed genes (Xu et al., 2021). The reprocessed dataset was composed of 29 clusters which were annotated to 10 identities using known marker genes (Supplemental Figure 9A). The limited available cell-type specific markers for “meristem base” and “meristem boundary” did not allow to differentiate these two cell-types. Similarly, we could not distinguish specific clusters for epidermis, but four clusters (clusters 7, 9, 20, 28) were annotated as “epidermis, determinate lateral organ”, as marker genes for these two cell-types were highly expressed. No clear signal was identified for “bundle sheath” marker genes, and one cluster (cluster 19), lacking specific expression of any of the available marker genes, was labeled “unclear” (Supplemental Figure 9A).

An initial MINI-EX run using default parameters (see Methods) returned a total of 842 regulons describing the interaction of 196 TFs within 28 clusters, corresponding to 14.7% (196/1332) of the expressed TFs, a considerably lower fraction compared to Arabidopsis and rice. The transition from the first step of MINI-EX to the last is accompanied by a large decrease in the number of analyzed TFs (Supplemental Figure 8B). While for rice and Arabidopsis this decrease is rather gradual, in maize we observed a drastic reduction characterizing only the last step (from 1249/1267 TFs from the TFBS-enriched regulons to 196 TFs in the final regulons - Supplemental Figure 8B). To test whether a more relaxed filter would mitigate this effect, an additional run of MINI-EX was performed setting the filter on the percentage of cells within the cluster expressing the TF to the minimum (step 3b - the TF needs to be expressed in the cell cluster, regardless of the percentage of cells). MINI-EX returned 6150 regulons for a total of 1148 TFs, including 90% of the expressed TFs (Supplemental Figure 8B, faded green, Supplemental Dataset 5). To evaluate the newly inferred regulons, the expression specificity of the top 150 prioritized regulons was analyzed (Supplemental Figure 9B). In contrast to Arabidopsis, only few maize TFs have an experimentally validated function. However, annotations transferred using orthology showed the top-prioritized regulators to specifically control the same cell-types as their respective Arabidopsis orthologs. Notable examples are VND4, VND7, TMO5, TMO5-like genes in xylem clusters (Kubo et al., 2005; Smet et al., 2019), OPB2, PEAR2, APL, DOF5.6 in phloem clusters (Bonke et al., 2003; Skirycz et al., 2006; Miyashima et al., 2019), and Zm00001d018930 and Zm00001d031009, orthologs of the Arabidopsis TCX2/SOL2 and TCX3/SOL1 genes, known to be involved in the regulation of cell division (Simmons et al., 2019), controlling regulons in cell cycle related clusters (Supplemental Figure 9B). Interestingly, an ortholog of Arabidopsis BZO2H2/bZIP9 was also found among the top phloem regulators in maize, further confirming the importance of this TF regulating gene expression in this specific tissue.

Next, for well characterized cell-type specific regulators (Xu et al., 2021) the percentage of cells expressing the TF for each inferred regulon was plotted (Figure 5A). Among the broadly expressed regulators we found KN1, known to be a core TF of meristematic tissue, controlling regulons in several meristem clusters, but also in vascular tissues, as previously reported (Jackson et al., 1994; Xu et al., 2021). Regulons for ZmCUC3-LIKE, known to control the initiation of axillary meristems in Arabidopsis (Li et al., 2020; Xu et al., 2021), and ZmYAB14, marker for determinate lateral organs (Strable et al., 2017), were inferred for all adaxial meristem periphery and determinate lateral organs clusters, respectively, and in some cell-cycle related clusters (Figure 5A). Similarly, vascular tissue regulons, containing regulators such as ZmTMO5 and TMO5-likes, and ZmAPL, were inferred for xylem and phloem clusters, respectively (Figure 5A). Remarkably, relaxing the filter on TF expression (step 3b) allowed to include relevant regulons which showed a highly specific expression among the cells of a cluster, such as ZmPEAR1-2/LIKE in the phloem cluster, and ZmHDZIV6 in epidermis (Figure 5A) (Javelle et al., 2011; Xu et al., 2021).

**Figure 5.**
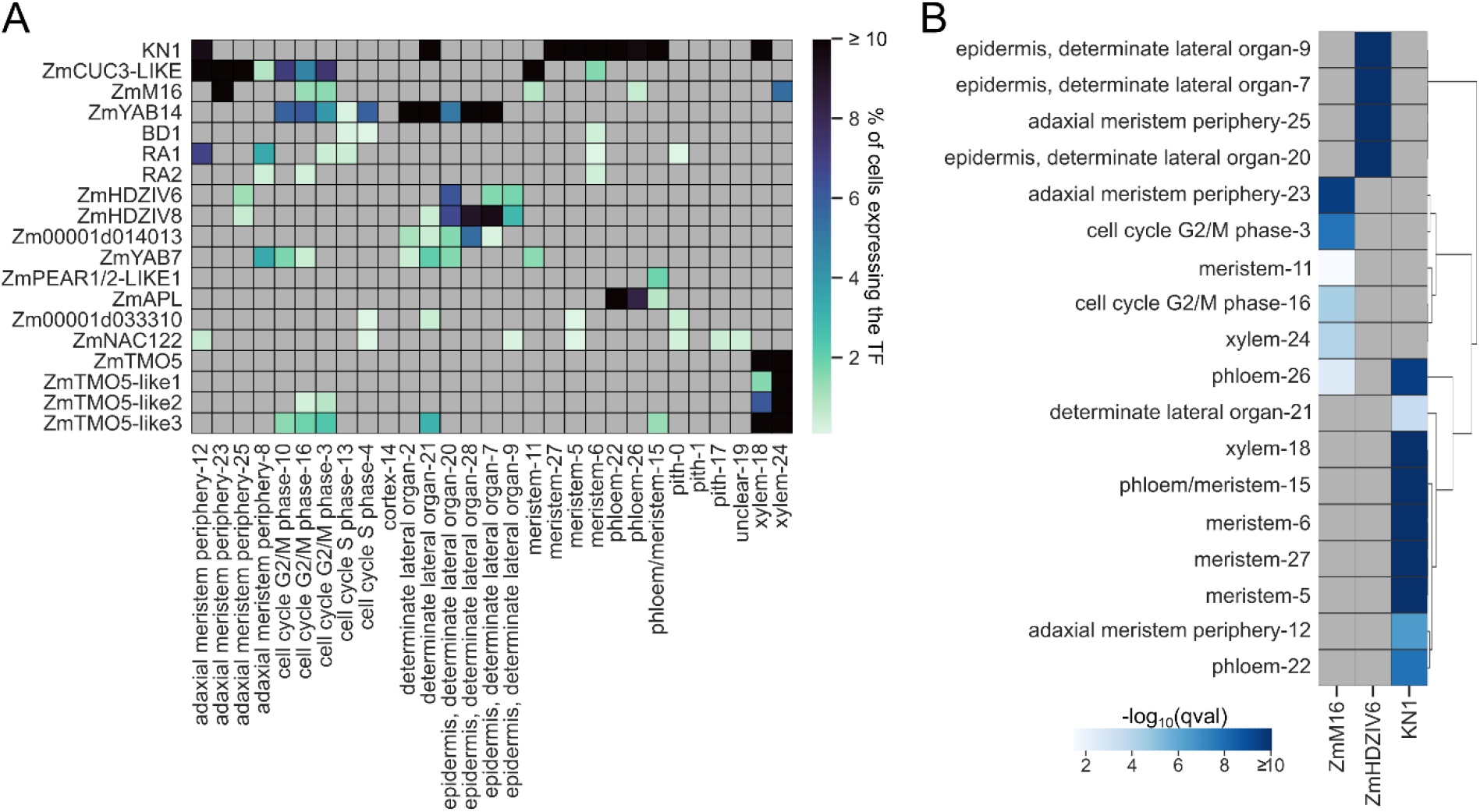
Expression and validation of cell-type specific regulators of maize developing ear. (A) Heatmap showing the percentage of cells expressing the TF in the cell cluster the regulon was inferred for. (B) Clustermap showing the significance of overlap (hypergeometric test, FDR corrected p-value < 0.05) between experimentally validated TGs from bulk ChIP-Seq and the predicted TGs in the different single-cell clusters for KN1, ZmHDZIV6 and ZmM16 regulons. Gray boxes indicate single-cell clusters for which no KN1, ZmHDZIV6 or ZmM16 regulon have been inferred.

To validate the predicted interactions data from bulk ChIP-Seq experiments was collected for three TFs: KN1, ZmM16 and ZmHDZIV6 (Bolduc et al., 2012; Xu et al., 2021). The predicted TGs for each of the MINI-EX inferred regulons related to the three TFs, were compared to their experimentally validated TGs (see Methods), and the significance of their overlap was tested. All predicted regulons showed a significant overlap with the experimentally validated TGs, with a higher significance for the cell-types the TF is known to act in (Figure 5B, hypergeometric test, FDR corrected p-value < 0.05). For each TF, functional enrichment of the regulon’s TGs was performed per cell-type (Supplemental Figure 9C). In line with their function and the cell-type they were inferred in, TGs for KN1 regulons in *meristem-27* and *6* were enriched for “auxin homeostasis” (Bolduc et al., 2012), while its TGs in *xylem-18* were enriched for “xylem development” (Townsley et al., 2013). Similarly ZmM16’s TGs in *adaxial meristem periphery-23* were enriched for “sepal development” and “specification of floral organ identity” (Xu et al., 2021), and ZmHDZIV6 regulons for “epidermal cell fate specification”, “cuticle development” and “wax biosynthetic process” in all epidermal clusters (Supplemental Figure 9C).

Taken together these results highlight how MINI-EX successfully identifies relevant cell-type specific regulators for maize and how MINI-EX parameters can be tuned to properly accommodate specific datasets or users’ needs.

## Discussion

The advent of multicellularity is one of the major innovations characterizing eukaryotic evolution. This was accompanied by the acquisition of differentiation programs allowing every cell to acquire a specific identity, specialize its function, and consequently resulting in an improved species fitness. As the genomic information is identical for all cells of an organism, what defines the identity of a cell is its transcriptome, often referred to as the cell fingerprint (Arendt et al., 2016). To untangle cell-type specific expression regulation we developed MINI-EX, a motif-informed GRN inference method which uses single-cell expression data to infer high-quality networks controlling the organization of different cell-types in plants. MINI-EX combines the power of expression-based network inference and TFBS enrichment to identify regulatory relationships among TFs and TGs. Single-cell cluster-specific genes are leveraged to assign cell-type specificity to each of the inferred regulons, which are further filtered for TF’s expression, on the assumption that a TF must be expressed in a number of cells within the cluster to be regulatory active. We showed that filtering regulons by TF expression, rather than upregulation, increased the F1 score from 13.6% to 21%, together with an increased precision and recall (from 14.7% to 15.4%, and from 12.6% to 33%, respectively). MINI-EX provides a precise and customizable approach to prioritize relevant regulons based on network centrality measures, expression specificity and functional relevance of TGs. The default settings of MINI-EX have been chosen by extensively testing different parameters using the Arabidopsis Interactions-Gold Standard enriched for root-interactions. However, as optimal parameters might vary for different species, as demonstrated for maize, these are easily adjustable according to the user’s requirements.

We compared the performance of our method with other state-of-the-art methods for single-cell GRN inference, GRNBoost2 and SCENIC (Aibar et al., 2017; Moerman et al., 2019), and showed that MINI-EX outperforms both methods. The regulons inferred by MINI-EX showed a higher F1 for 72/82 and 12/18 of the commonly inferred regulons for GRNBoost2 and SCENIC, respectively, demonstrating the recovery of a higher number of known interactions. MINI-EX’s performance was evaluated using two different gold standards, focused on known interactions and biological gene function. While our evaluation using the Interactions-Gold Standard reported an overall precision of 15.4%, we showed that 23.5% of the top 150 regulons were related to TFs known to be involved in root development (“relevant TF”, Functional-Gold Standard), indicating that the actual precision is probably higher. It is worth noticing that the lower precision obtained using the Interactions-Gold Standard is most likely due to the different regulon sizes in the two networks (median of 42 TGs in the Interactions-Gold Standard vs 510 TGs in MINI-EX). The Interactions-Gold Standard was constructed by combining different data types and is incomplete, which means that many novel correct interactions are labeled as false positives, making it more difficult to obtain high precision values compared to high recall. The increased performance of MINI-EX is primarily due to the use of TF-family information, together with an ensemble approach during TFBS enrichment analysis. For the latter, it was shown that FIMO and Cluster-Buster, two frequently used motif mapping tools in plants (Frith et al., 2003; Grant et al., 2011) produce complementary results when evaluating TFBS mappings using ChIP-Seq in Arabidopsis (Kulkarni et al., 2019).

We demonstrated that for the analyzed Arabidopsis scRNA-Seq dataset imputation did not to lead to a strong improvement of the inferred GRN. The lower performance observed could be affected by the normalization performed during the imputation step, as it is strongly recommended when running GRNBoost2, to avoid using normalized counts and sample normalized matrices because they might influence the co-expression analysis (Crow et al., 2016). In an extensive analysis, Hou and colleagues (Hou et al., 2020) revealed how imputation of missing data improved several steps of single-cell data analysis, such as clustering and trajectory analysis, however, no effect on GRN inference was studied. Ly and Vingron (Ly and Vingron, 2022) demonstrated that imputation methods are dataset-dependent. As such, a possible explanation for the poor performance can be attributed to the fact that the processed Arabidopsis dataset is of high-quality, thus, we cannot exclude that results would be different for GRNs inferred from lower-quality single-cell input data.

Through the integrative prioritization strategy based on weighted Borda counts, MINI-EX identified half of the expected root-related TF (“relevant TF”) within the top 37.8% of regulons, 285 ranks earlier than the best-performing single metric (cluster-specificity). MINI-EX successfully identified relevant root TFs as well as recently characterized ones, such as ERF13, associated to lateral root development (Dossa et al., 2020), regulating procambium and pericycle, and BRAVO/MYB56, described as a modulator of cellular quiescence (Betegón-Putze et al., 2021), controlling *QC-37*.Moreover, overlapping the inferred regulons with genes showing an association with root phenotypes revealed how unknown genes, representing promising candidates for functional characterization studies, were prioritized well. Examples include genes known to be involved in root morphogenesis such as KNAT7 and HAT4 (Steindler et al., 1999; Zhong et al., 2008), but more interestingly, also genes with unknown functions such as AT2G03470 in *xylem-31*, WRKY3 in *trichoblast-27*, and bZIP6 in several stele-related clusters. Notably, the root-GWAS reported phenotype associated to bZIP6 is related to lateral root density (Julkowska et al., 2017), in line with the fact that it appears as main regulators of several pericycle clusters. Other relevant examples are represented by genes which have already an experimental validated function such as WRKY26, known to be involved in thermotolerance (Li et al., 2011), and ZAT12, active in abiotic stress signaling (Davletova et al., 2005), both regulating several LRC clusters. While the initial expression-based network, inferred using GRNBoost2, offers a solid basis for MINI-EX’s final regulons, the inherent bias towards positively correlated TF - TG gene pairs (mean Pearson correlation coefficient between TF - TG of 0.18) makes it difficult to reliably label the predicted regulators as activators or repressors. As the same TF can have a dual function depending on the cellular context (bifunctional TFs), *in vivo* functional validation is most likely needed to characterize the regulatory role of specific regulators in individual cell types.

The applicability of MINI-EX on different plant species was further demonstrated on rice and maize datasets. An in-depth analysis showed that, for all three species, the TFBS enrichment step did not lead to a large loss of regulons, indicating that our motif collection is reasonably complete. In the analyzed rice single-cell dataset, TFs tend to be expressed throughout different cells, but not consistently across cells of the same type. This pattern was even more pronounced in the maize ear dataset, where the application of a more relaxed filter for TF expression alleviated this problem and resulted in the recovery of additional know cell-type specific regulators. Finally, we showed how MINI-EX can be leveraged to dissect regulatory information from bulk experiments across the different rice root cell-types. By overlapping target genes previously identified by bulk ChIP- and RNA-Seq on overexpression lines of three rice NAC TFs (Chung et al., 2018) with the cell-type specific regulons inferred by MINI-EX, we could resolve specific cell-type regulation better explaining the observed phenotypes.

We presented MINI-EX, an efficient method for GRN inference for scRNA-Seq data in plants. The extensive validations and obtained results described here showed how MINI-EX represents a powerful tool to study gene regulation at single-cell level and how it accurately prioritizes regulons through an integrative prioritization strategy leveraging network characteristics, expression specificity and functional relevance. Furthermore, we presented different examples indicating how it can help in hypothesis generations, not only in Arabidopsis, but also in economically relevant species such as cereals, and comparative studies.

## Methods

### Motif collection and mapping

TF motifs modelled as position weight matrices (PWMs) for Arabidopsis, rice and maize were collected from the CisBP 2.00 (downloaded on Dec 2019) (Lambert et al., 2019) and JASPAR2020 (Fornes et al., 2020) databases. For Arabidopsis these were combined with a collection of manually curated motifs (Kulkarni et al., 2018). The lists of TFs used was further extended by adding TFs belonging to families represented by at least one motif in our motif collection. TF-families were assigned to TFs based on PlantTFDB (Jin et al., 2017) and PlntTFDB (Riaño-Pachón et al., 2007), yielding in 57, 42, 41 TF families for Arabidopsis, rice and maize, respectively. PlantTFDB was used as a primary reference for TF-family assignments. A total of 1699, 1244, 1335 motifs, corresponding to 1117, 724, 1234 TFs for Arabidopsis, rice and maize, respectively, were collected. The TF-family extension increased the number of TFs to 1877, 1643, 2148 TFs, respectively. TF motifs were compared in a pairwise manner using RSAT-compare matrix (Medina-Rivera et al., 2015) and motifs with a normalize correlation (Ncor) equal to 1 were considered duplicates and removed from the collection. TF motifs were mapped to a genes’ regulatory regions using Cluster Buster (version compiled on Sept 2017)(Frith et al., 2003) and FIMO (meme version 4.11.3)(Grant et al., 2011). Cluster Buster was run with the -c parameter set to 0 and PWMs were scaled to 0-100. FIMO was run with default parameters and PWMs were scaled to 0-1. The regulatory regions used for motif mapping were extracted from Arabidopsis Araport11 (Cheng et al., 2017), Rice JGI v7.0 (Ouyang et al., 2007) and Maize AGP v4.0 (Jiao et al., 2017) gff3 files, only retaining the longest splicing variants, downloaded from PLAZA Dicots 4.5 (Van Bel et al., 2018). These regions were initially defined as 5000 bp upstream and 1000 bp downstream from translation start/stop sites. Introns were included in the regulatory regions, while coding exons were excluded. Importantly, if another gene was present within the 5000 bp-1000 bp window, this region was cut where the other gene started. For Arabidopsis, this resulted in most upstream regions being much smaller than 5000bp, with a median length of 1489 bp and mean of 2022 bp. The optimal number of matches per tool and motif were calculated similarly to Kulkarni et al., 2019 using a collection of ChIP-Seq peaks intersected with our promoter definition for 40 and 68 TFs (Supplemental Table 2) for Arabidopsis and maize, respectively. This resulted in the selection of the top 4000 and 8000 matches from Cluster Buster and top 7000 and 16,000 matches from FIMO, for Arabidopsis and maize, respectively. For rice, where we lacked a high-quality ChIP-Seq data to perform a similar calibration, a proportional number of matches relative to the number of total genes was chosen, resulting in top 6000 and top 10,000 matches for Cluster Buster and FIMO, respectively. These matches were combined to create an Ensemble motif mapping file (motif-TG) used for the TFBS enrichment in MINI-EX.

### Single-cell RNA-Seq data processing

Raw Arabidopsis, rice and maize fastq files were downloaded from the Sequence Read Archive (SRA) (SRP235541, SRP173393, SRP292306, SRX7814225, SRR12276825, SRR12276824, SRR12276805) and processed using Cell Ranger v3.1 (10X Genomics). When replicates were present they were aggregated using the Cell Ranger function *aggr*. The reference was built using the Cell Ranger function *mkref* with gtf file filtered for protein coding genes. For Arabidopsis and maize the genome and gtf files were downloaded from Ensembl-Plants (TAIR 10.45 and AGP v4.0, respectively), while for rice genome and gff3 files were downloaded from PLAZA Dicots 4.5 (Ouyang et al., 2007; Van Bel et al., 2018). The gff3 file was converted to gtf using *gffread* command from cufflinks (Trapnell et al., 2010). All gtf files were filtered to only keep the longest splicing variant.

The filtered gene-to-cell matrices were further processed using Seurat v3.1.2 (Stuart et al., 2019). Specifically, the filtered feature-barcode matrix output of Cell Ranger was used to create a Seurat Object and cells which showed too few genes (< 200) and too high UMI counts (>100,000 for Arabidopsis, and > 10,000 for rice and maize, respectively) were filtered out. These filters were chosen upon QC metrics visualization using the *FeatureScatter* function of Seurat plotting “nCountRNA” (UMI counts) versus “nFeature_RNA” (gene counts) to identify outliers. The datasets were normalized using the *NormalizeData* function with normalization.method=“LogNormalize”, scaled and centered regressing out variance related to UMI counts using the *ScaleData* function with vars.to.regress =“nCount_RNA”, and highly variable genes were identified using the Seurat function *FindVariableFeatures*. These were used for PCA dimensionality reduction, where 50 PCs were computed and used for downstream processing. The K-nearest neighbor (KNN) graph was constructed using the Seurat function *FindNeighbors* with dims=1:50, and using the function *FindClusters* with a resolution of 2 for Arabidopsis (Wendrich et al., 2020) and maize and 1 for rice and the other Arabidopsis datasets (Denyer et al., 2019; Kim et al., 2021), 41, 20, 18, 19 and 29 clusters were identified for Arabidopsis (Wendrich et al., 2020; Denyer et al., 2019; Kim et al., 2021), rice and maize, respectively. Marker genes for each cluster were selected using the function *FindAllMarkers* with *only.pos* set to TRUE to only retrieve upregulated genes.

For Arabidopsis, cluster identities were assigned based on the expression of known cell-type marker genes (Wendrich et al., 2020) (Supplemental Figure 4A), and by mapping the top 20 upregulated genes for each cluster to known cell-type specific datasets (Li et al., 2016). For rice and maize, cluster identities were assigned based on the expression of known cell-type marker genes (Xu et al., 2021; Liu et al., 2021). The Arabidopsis published and processed dataset (Wendrich et al., 2020) was kindly provided by the authors. The Jaccard Index used to compare the regulons inferred from the original and reprocessed dataset was calculated as the intersection over the union of the top 50 regulons per cell-type/cluster.

### Construction of Gold Standards

The Interactions-Gold Standard used to evaluate MINI-EX parameters and performance was constructed starting from protein-DNA interactions obtained from ChIP-Seq data, Y1H data and literature curated data (Supplemental Table 3). NarrowPeaks were downloaded from PCBase (Chow et al., 2019), with the exception of peaks from Song et al., 2016. Peaks were annotated to the closest gene. Only target genes confirmed by at least two replicates were retained. NarrowPeaks for 26 TFs were downloaded from GEO (GSE80564) (Song et al., 2016) and annotated to the closest gene. As replicates were not reported but only peaks corresponding to ABA-treated and mock samples, we retained only TGs confirmed by both samples. For other TFs, when narrowPeaks were available, these were downloaded and annotated in a similar manner, otherwise, lists of high confidence target genes were retrieved from supplementary material of corresponding papers (Supplemental Table 3).

To make the Interactions-Gold Standard root-specific, it was first filtered to only keep TFs expressed in the single-cell expression atlas, and afterwards the TGs of each TF that were enriched for the upregulated genes of the root single-cell dataset were retained (hypergeometric test, FDR corrected p-value < 0.05). Only TF with at least 10 reported interactions were retained. The resulting Interactions-Gold Standard reports 82,303 interactions for 115 TFs, with a median of 42 TGs per TF.

The Functional-Gold Standard was constructed based on Gene Ontology annotations for biological process (BP) (downloaded on September 2020 from http://geneontology.org/gene-associations/tair.gaf.gz). All genes with experimental evidence (GO evidence code: EXP, IMP, IDA, IPI, IGI, IEP) or manually curated (GO evidence code: TAS, NAS, IC) GO annotation related to root biology (searching for “root”, “xylem”, “phloem”, “vascular”, “trichoblast”, “trichome”, “vasculature”, “stele”, “tracheary”, “procambium”, “sieve”) were selected, retaining in total 845 Arabidopsis genes. Functional annotation used for the leaf dataset was obtained by combining the experimentally/manually curated GO annotation related to leaf biology (searching for “leaf”, “xylem”, “phloem”, “vascular”, “bundle sheath”, “chloroplast”, “photosynthesis”, “photosystem”, “vasculature”, “vascular”, “stele”, “tracheary”, “procambium”, “sieve”) with a literature curated list of relevant leaf TFs (Supplemental Dataset 3).

### MINI-EX network inference

In the first step GRNBoost2 (Moerman et al., 2019) is used to obtain the expression-based network using as inputs a list of TFs and the gene-to-cell matrix filtered for expressed genes (read count >1). Next, each expression-based regulon is tested for enrichment of the TFBSs relative to TF motif(s) recognized by the TF controlling the regulon, using a hypergeometric test and the Ensemble motif mapping file. Only regulons showing a significant enrichment (FDR corrected p-value < 0.001) for a motif directly recognized by the TF controlling the regulon or for a motif recognized by other TFs belonging to the same TF family are retained. Finally, the TFBS-enriched regulons are tested for the enrichment of single-cell cluster-specific upregulated genes. Only regulons with a significant enrichment (hypergeometric test, FDR corrected p-value < 0.001) and for which the TF is expressed in at least 10% of the cell of the cluster are retained.

Optimal thresholds for each step were determined upon calculation of precision, recall and F1 score of multiple cutoffs (Supplemental Figure 1). Precision was defined as the number of TF-TG interactions confirmed by the Interactions-Gold Standard divided by the total number of inferred TF-TG interactions, while recall was calculated by dividing the number of TF-TG interactions confirmed by the Interactions-Gold Standard with the total number of TF-TG interactions reported by the Interactions-Gold Standard. For the calculations the gold standard was filtered to only include TFs with inferred interactions and the inferred network was filtered to only retain interactions related to TFs part of the gold standard. As a result, all predictions of GRNBoost2 (top_100% - step 1), top700 marker genes (top700_cellClusters – step 3a), and 10% expression filtering for the regulon’s TF (EXPfiltered_10 – step 3b) were chosen as optimal parameters (Supplemental Figure 1). For maize, the TF expression filtering (step 3b) was adjusted to >0%.

MINI-EX provides basic network centrality measures for each regulon calculated using the python package networkx (https://networkx.org/), where each centrality measure is normalized by network size, and functional enrichment based on experimentally validated and/or manually curated GO terms (hypergeometric test, FDR corrected p-value < 0.05). To prioritize regulons for their importance in the regulation of a cell cluster, MINI-EX uses the cluster-specificity (FDR corrected p-value < 0.001 for cell cluster enrichment), together with the network centrality measures, to rank each regulon based on single metrics. Successively, it calculates the geometric mean of the individual ranks to obtain the Borda rank. In case a list of expected GO term is provided, MINI-EX also includes a functional-specificity measure (FDR corrected p-value < 0.05 for functional enrichment of the relevant GO terms among the regulon’s TGs) within the Borda calculations, and progressively evaluates all combinations of metrics based on the retrieval of half of the relevant regulons earliest in the rank (R50). In this latter case, each metric is weighted by its importance (ranki/R50) before being combined for Borda ranks.

The MINI-EX pipeline has been implemented in the Nextflow workflow (DLS2) management system and uses a specific Singularity container for all necessary dependencies. A detailed illustration of the pipeline is described in the public GitHub repository https://github.com/VIB-PSB/MINI-EX. The default parameters used by MINI-EX can be easily changed by the user in the Nextflow config file to best accommodate specific requirements.

### Robustness of MINI-EX single-cell network inference and comparison with other methods

To evaluate the stability of MINI-EX the inferred regulons were compared to regulons obtained from down-sized expression matrices. Varying fractions (from 10% to 90% of the initial cells) of the initial matrix were randomly sampled for 10 times and used as inputs of MINI-EX. Similarly, to test the robustness of MINI-EX towards missing values, MAGIC and SAVER imputation methods (Huang et al., 2018b; van Dijk et al., 2018) were run on the initial expression matrix with default parameters and the imputed matrices were used as input for MINI-EX. To compare MINI-EX with SCENIC (Aibar et al., 2017), we adapted the SCENIC pipeline to Arabidopsis data. First, a genome ranking file was constructed, where, for each motif, the Arabidopsis genes were ranked based on the Cluster Buster (Frith et al., 2003) scores as recommended by the authors. Then, a TF information file linking each TF to the recognized TF motifs was built and used, together with the gene-to-cell expression matrix, as inputs for the SCENIC pipeline. For the comparison with GRNBoost2 (Moerman et al., 2019), we compared the results of the first step of MINI-EX with the final MINI-EX regulons.

### GWAS evaluation

GWAS association data was collected from AraGWAS (downloaded on Nov 2019)(Togninalli et al., 2018) and screened for studies associated to root phenotypes. All significant associations (Bonferroni corrected p-value < 0.001) with a Minor Allele Count (MAC) > 5 were retained. Only SNPs on coding sequences were kept, resulting in 36 studies and 560 genes. As MINI-EX uses the upregulated genes of each cell cluster to filter regulons for cell-type specificity, only the 187 root-GWAS genes which appeared as upregulated in at least one cell cluster (named “upregulated root-GWAS”) were considered when studying the GWAS representation within regulon’s TGs.

### Rice bulk dataset collection

Rice NAC RNA-Seq and ChIP-Seq bulk data were collected from Gene Expression Omnibus database (GSE102921) (Chung et al., 2018). Only data relative to OsNAC5, 6 and 9 was used, as no OsNAC10 regulon was inferred by MINI-EX. Processed annotated peaks were downloaded and rice ID were converted from IRGSP 1.0 to MSU 7. As no processed DEG data for the single TFs was available, raw RNA-Seq fastq files were downloaded and processed with Curse (Vaneechoutte and Vandepoele, 2019) to obtain a gene-to-count matrix which was used for differential expression analysis using DESeq2 (Love et al., 2014). Genes which showed a log2 fold change > 1.5 or < −1.5, and an adjusted p-value < 0.05 were kept. Only TGs which resulted differentially expressed (up- and downregulated) and bound in the RNA-Seq and ChIP-Seq dataset, respectively, were considered as experimentally validated.

Cytoscape 3.7.2 (Shannon et al., 2003) was used for network visualization. Orthology information was obtained from PLAZA Dicots 4.5 (Van Bel et al., 2018).

### Maize bulk dataset collection

High confidence annotated peaks, defined as peaks in the two biological replicates whose midpoints showed a distance ≤ 300 bp, for ZmHDZIV6 and ZmM16 ChIP-Seq bulk data were collected from (Xu et al., 2021). KN1 modulated and bound genes, defined as genes annotated to high confidence peaks which showed differential expression (up- and downregulated) in *kn1-e1* loss-of-function mutant, were collected from (Bolduc et al., 2012). Gene IDs were converted from AGPv3.31 to AGPv4, and gene symbols were retrieved from PLAZA Arabidopsis’ orthologs and from the original publication (Van Bel et al., 2018; Xu et al., 2021). GO enrichment of KN1, ZmHDZIV6 and ZmM16 TGs was performed using a GO-gene file containing all annotations for biological process (BP) (downloaded from PLAZA monocots 4.5 (Van Bel et al., 2018), and the results were filtered to exclude too general parental terms.

## Data availability

MINI-EX’s source code and Nextflow DSL 2 pipeline is available in the following public repository https://github.com/VIB-PSB/MINI-EX under the GPL-3 license. An example with a reduced matrix and tutorial is also provided in the repository.

## Author contributions

KV and CF conceived the project and designed the research. NMP performed the TF motif mapping calibration for all the species and collected literature information for relevant leaf TFs. CF developed the method, CF and KV wrote the manuscript. All authors read and approved the final paper.

## Acknowledgments

We would like to thank Dr. Bert de Rybel, Dr. Moritz Nowack, Jim Renema and Dr. Tereza Vavrdova for the insightful discussions, Bert Droesbeke for the help with building the Docker image, Cecilia Sensalari and Dr. Svitlana Lukicheva for testing the pipeline, and Dr. Thomas Eekhout for providing the original processed Arabidopsis dataset. We would also like to thank Dr. Dries Vaneechoutte for his work on compiling the hypergeometric enricher used, Francois Bucchini for the initial parsing of the GWAS data, and Dr. Yvan Saeys for useful suggestions related to the imputation evaluation.

## Supplemental information

**Supplemental Figure 1.**
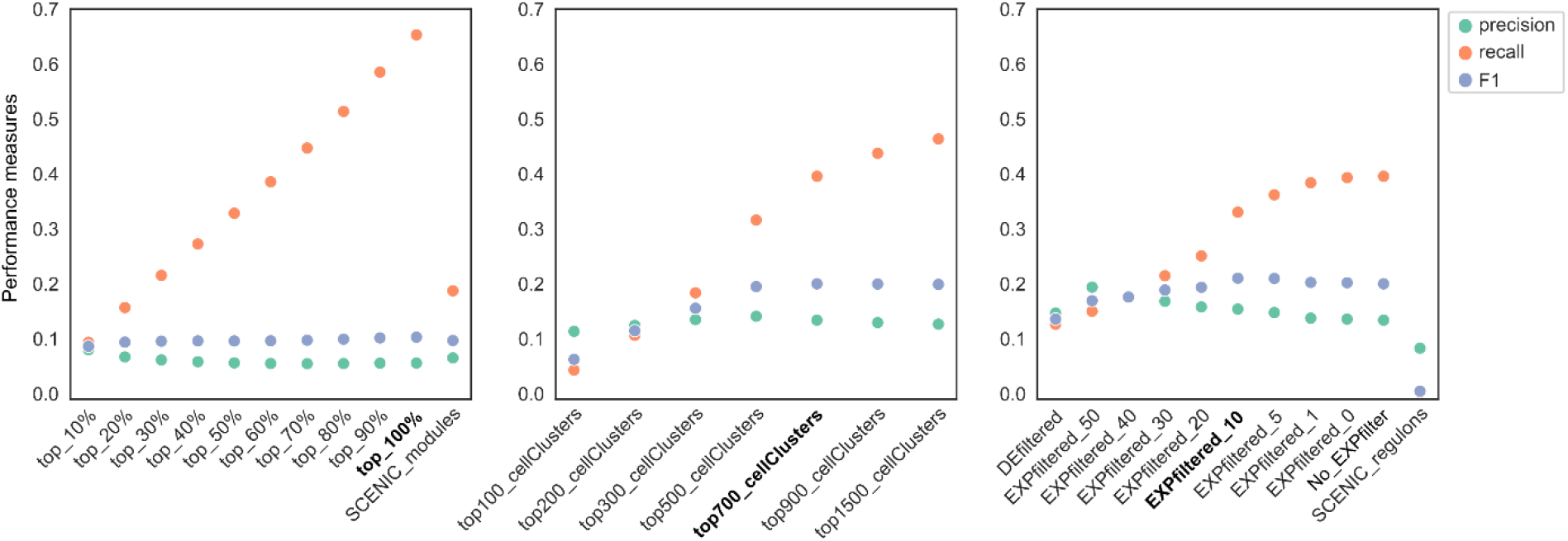
Benchmarking of optimal MINI-EX parameters. Point plot graphs indicating performance measures for different steps of the MINI-EX pipeline and SCENIC. Points indicate precision (green), recall (orange) and F1 score (violet). The parameters in bold are the ones showing the best performance and are used as default in MINI-EX.

**Supplemental Figure 2.**
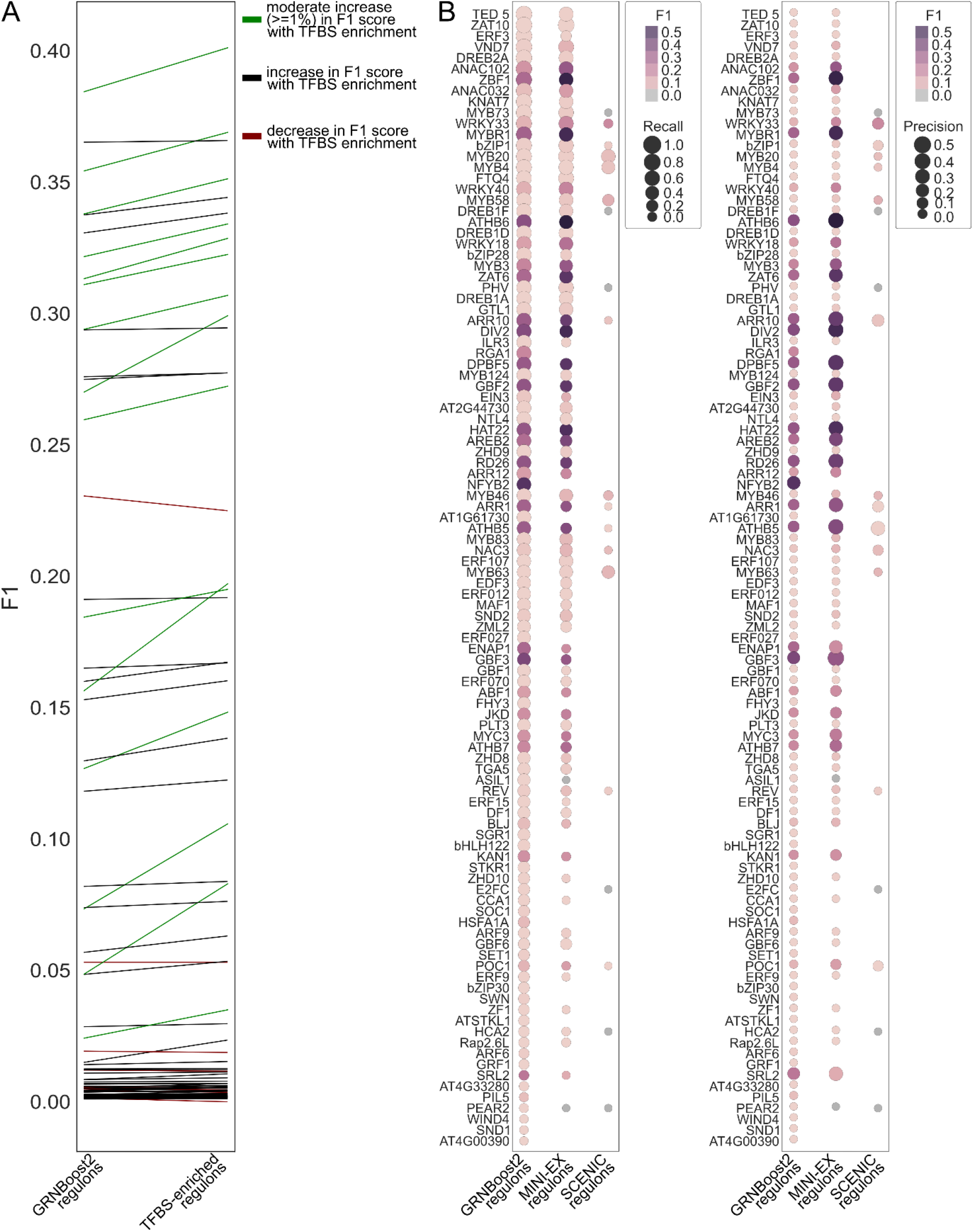
F1 score comparisons for MINI-EX and related methods. (A) Line plot reporting changes relative to F1 scores for 91 regulons belonging to the Interactions-Gold Standard that passed the TFBS enrichment step. Green lines indicate regulons which showed a moderate (≥ 1%) increase in F1, black lines indicate regulons which showed an increase in F1 score, and red lines indicate regulons which showed a decrease in F1 score after TFBS enrichment. The starting of the line indicates the F1 score for the GRNBoost2-inferred regulon, and the end of the line indicates the F1 score for the same regulon after TFBS enrichment. (B) Bubble plot comparing F1 scores of regulons inferred by GRNBoost2, MINI-EX and SCENIC. The size of the bubble indicates recall in the plot on the left and precision on the plot on the right, while the color represents F1 score.

**Supplemental Figure 3.**
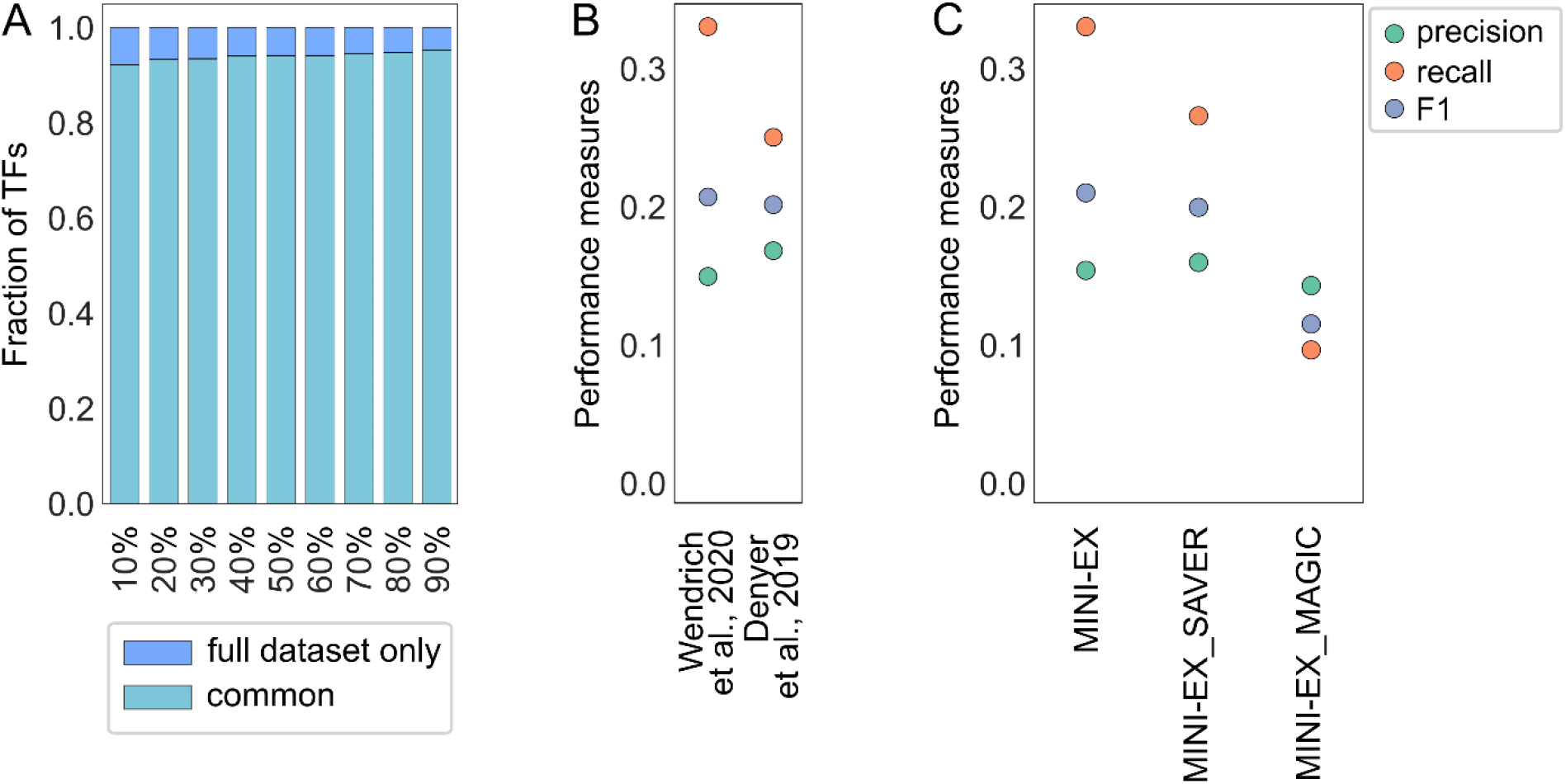
Stability of MINI-EX towards down-sampling and missing values. (A) Stacked bars indicating the fraction of TFs (n=696) from the original run identified by MINI-EX on the down-sampled matrices. (B) Point plot showing precision, recall and F1 score for the networks inferred from the two Arabidopsis’ root scRNA-seq datasets (Wendrich et al., 2020 and Denyer et al., 2019). (C) Point plot showing performance measures of MINI-EX run on the non-imputed matrix, and after data imputation using SAVER and MAGIC. Points indicate precision (green), recall (orange) and F1 score (violet).

**Supplemental Figure 4.**
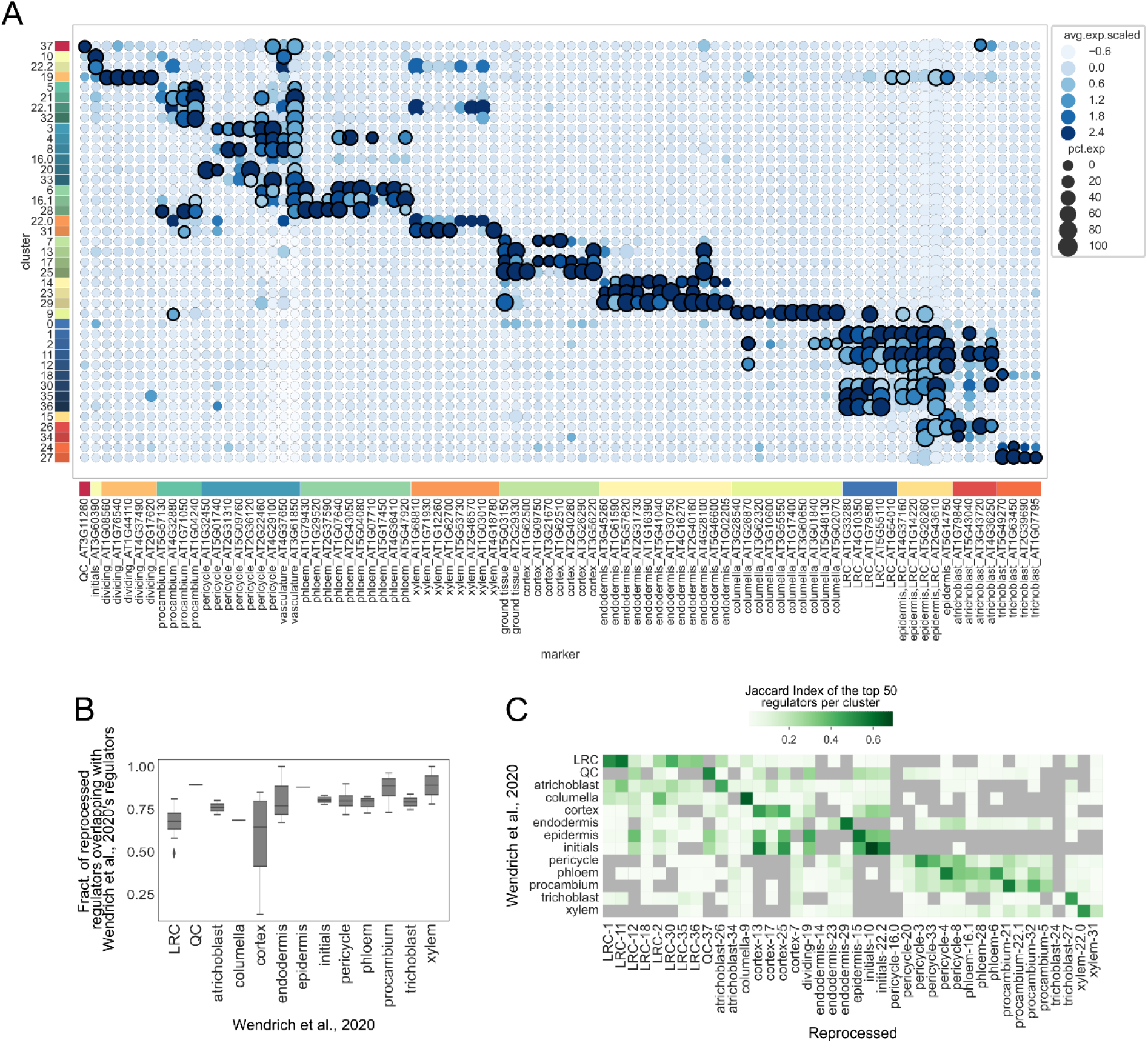
Reprocessing the Arabidopsis root dataset. (A) Bubble plot showing the expression of known marker genes from Wendrich et al., 2020 across the identified clusters. The color of the dots indicates the scaled average expression, while the size of the dots indicates the percentage of cells within the cluster expressing the gene. Dots with a thicker outline indicate genes which were identified as upregulated in the cell cluster. Marker genes and clusters are color-coded according to their annotation. (B) Box plots representing the fraction of regulators of the reprocessed dataset overlapping with the regulators inferred using the original dataset from Wendrich et al., 2020. (C) Heatmap showing the overlap (Jaccard Index) of the top 50 regulators between the original dataset (Wendrich et al., 2020) and the reprocessed one.

**Supplemental Figure 5.**
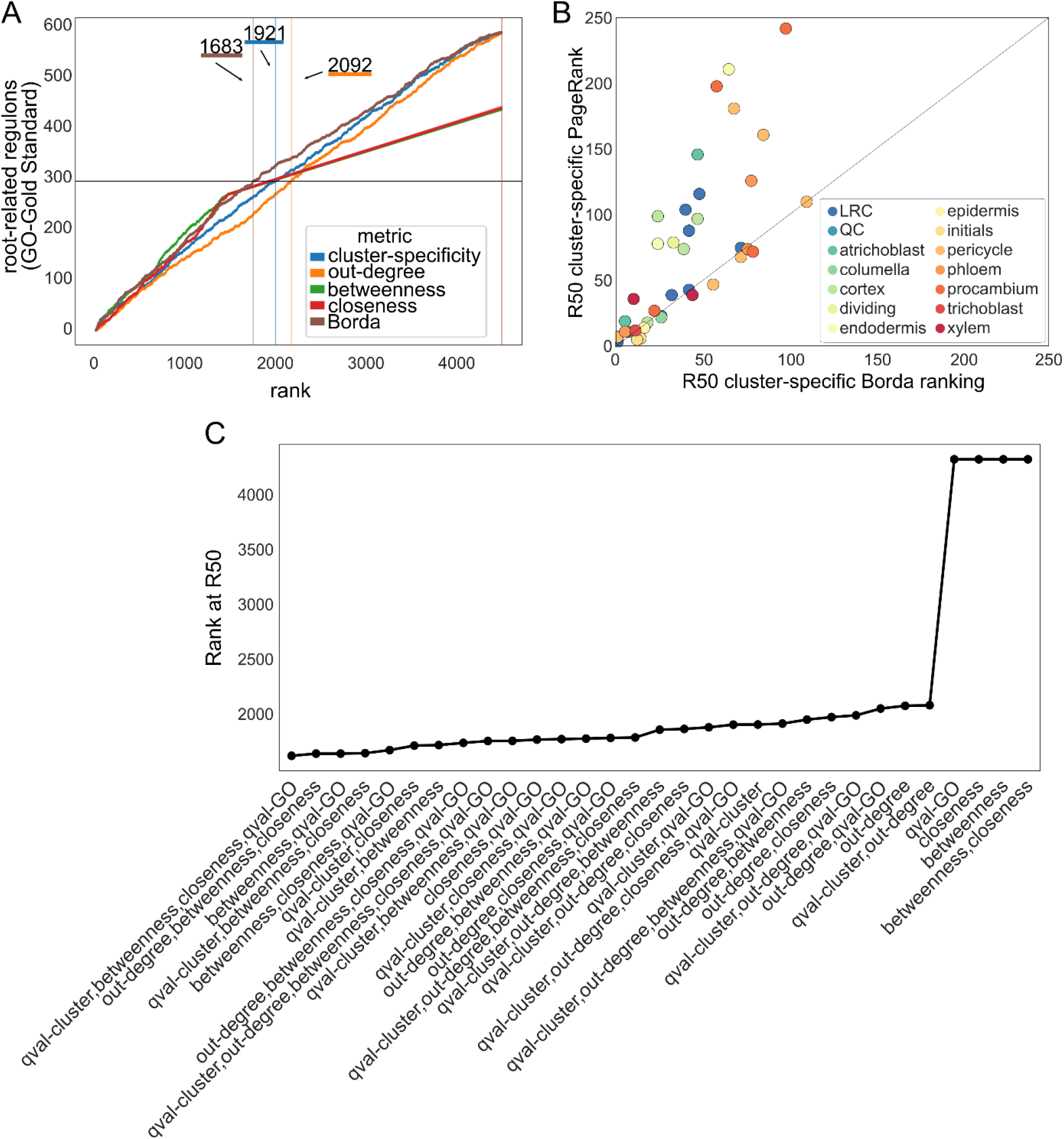
Evaluation of prioritization methods of relevant regulons. (A) All inferred regulons are ranked based on the four single metrics and Borda counts calculated on the geometric mean of the four metrics (x-axis), the identification of a root-related regulon (“relevant TFs”, from Functional-Gold Standard) is shown by the counts on the y-axis. The horizontal line indicates half of the root-related regxuulons (“relevant TFs”, from the Functional-Gold Standard), while the vertical lines indicate, for each metric, the rank at which 50% of the relevant root-related regulons is retrieved (R50). (B) Scatter plot indicating R50 values obtained from the cluster-specific Borda ranking (x-axis) and cluster-specific PageRank (y-axis) for each of the single-cell clusters. Dots are color-coded according to cell-types. (C) Point plot showing the rank at which half of the root-related regulons (from the Functional-Gold Standard) was retrieved for all possible combinations of the five metrics. “qval_cluster” and “qval_GO” refer to cluster-specificity and functional-specificity ranks, respectively.

**Supplemental Figure 6.**
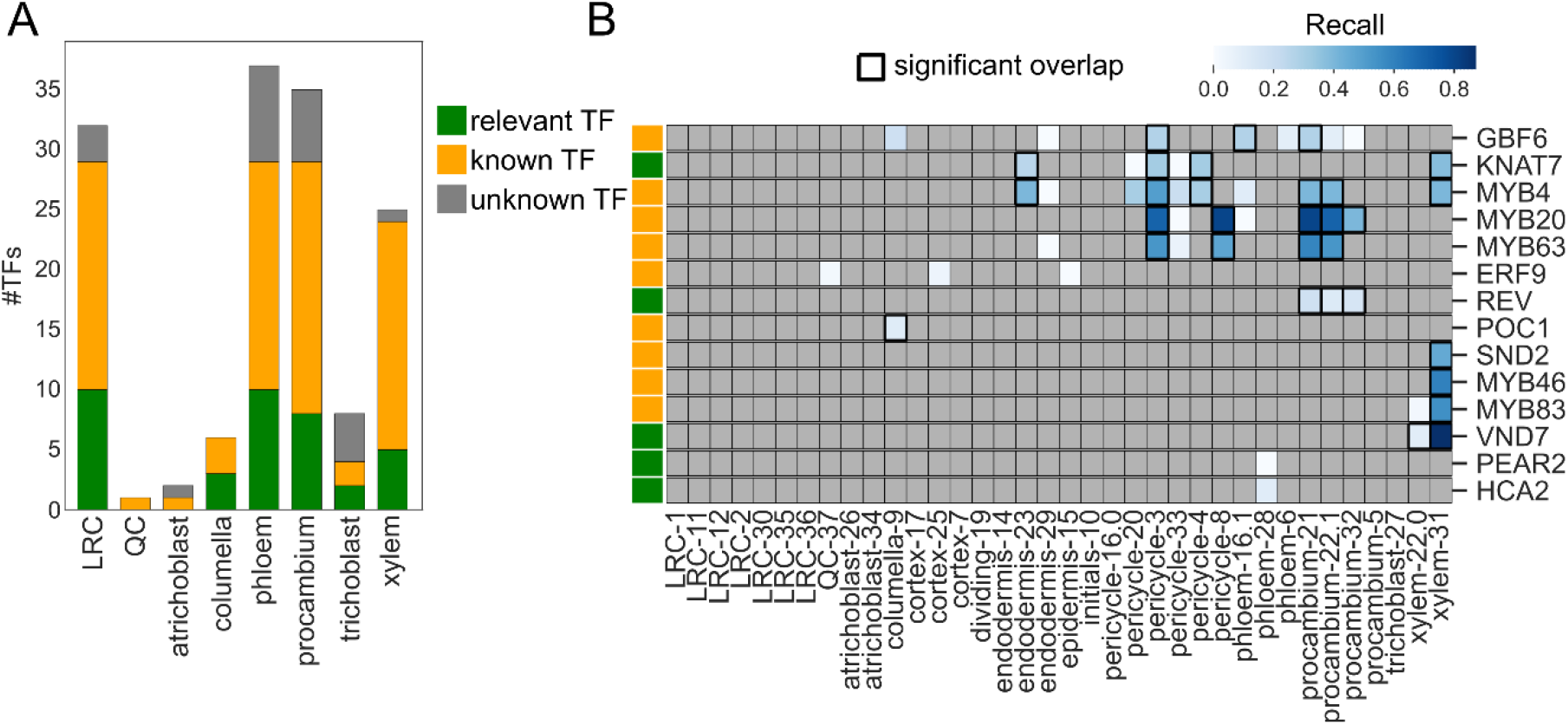
Evaluation of cell-type specific regulons and predicted target genes for 14 of the top 150 regulons. (A) Histogram showing the fraction of the top 150 regulons with a root-related GO annotation (from Functional-Gold Standard, “relevant TF”, green), another GO annotation (“known TF”, yellow), or unknown function (“unknown TF”) across different cell-types (B) Heatmap showing recall values for regulons controlled by 14 TFs part of the Interactions-Gold Standard. Cells with a thicker outline indicate a significant overlap (hypergeometric test, FDR corrected p-value < 0.05). Green and yellow boxes indicate whether the TF controlling the regulon has a root-related GO annotation (“Relevant TF”, from Functional-Gold Standard) or another GO annotation (“known TF”), respectively.

**Supplemental Figure 7.**
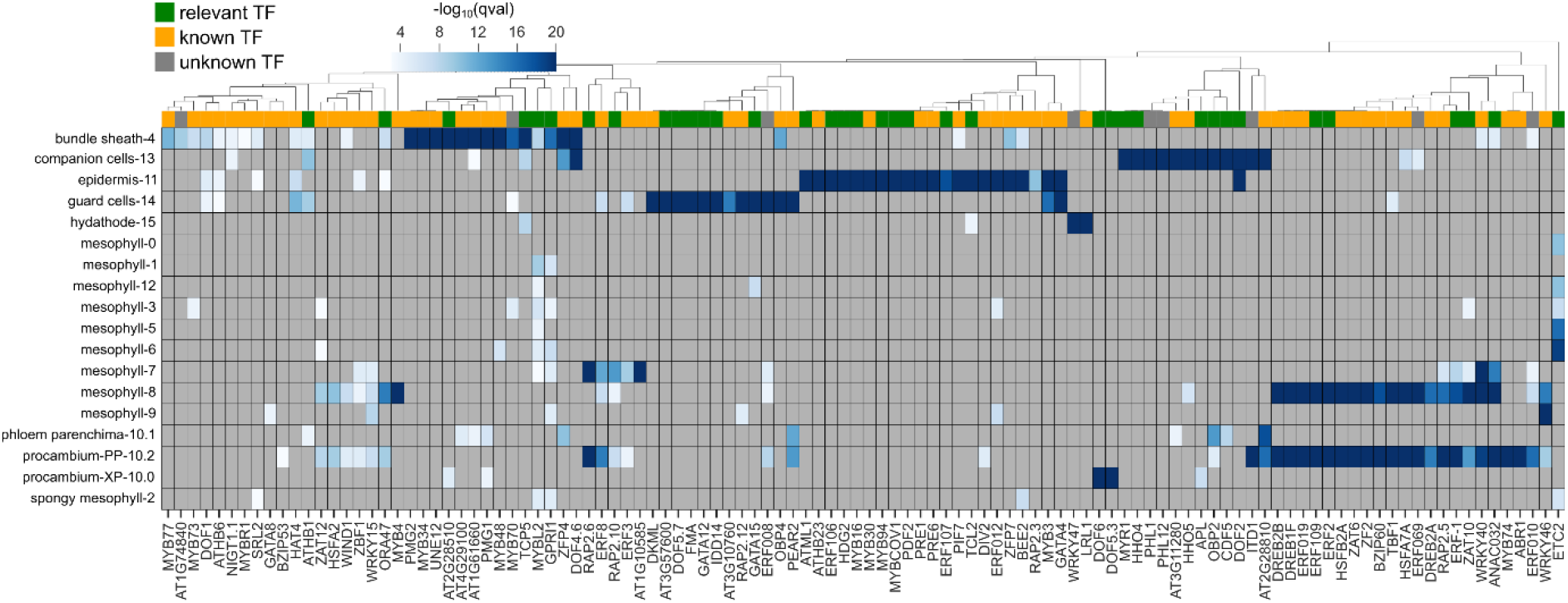
Expression specificity of the top 150 regulons controlling Arabidopsis leaf development. Clustermap showing cluster-specificity (the −log_10_(qval) of the single-cell cluster enrichment) of the top 150 inferred regulons. Green, yellow and gray boxes indicate whether the TF controlling the regulon has a leaf-related functional annotation (“relevant TF”), another functional annotation (“known TF”), or no functional annotation (“unknown TF”), respectively.

**Supplemental Figure 8.**
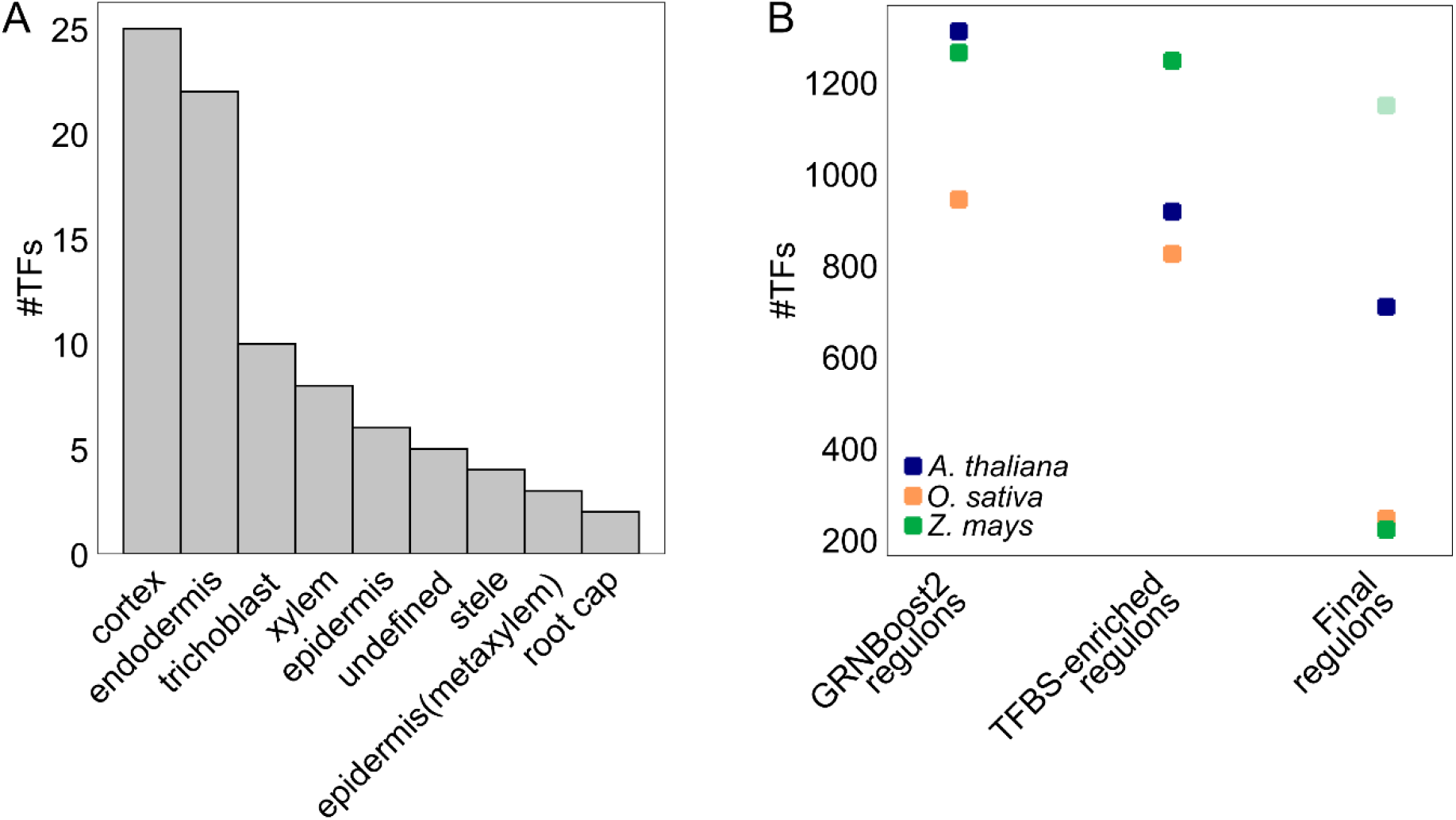
Properties of the GRNs inferred using MINI-EX. (A) Histogram showing the number of cell-type specific regulons per cell-type in the rice GRN. (B) Point plot showing the number of TFs retained after every step of MINI-EX, for Arabidopsis (blue), rice (orange) and maize (green). Faded green indicates the number of TFs retained by the last step of MINI-EX when the filter on TF expression is lowered in maize.

**Supplemental Figure 9.**
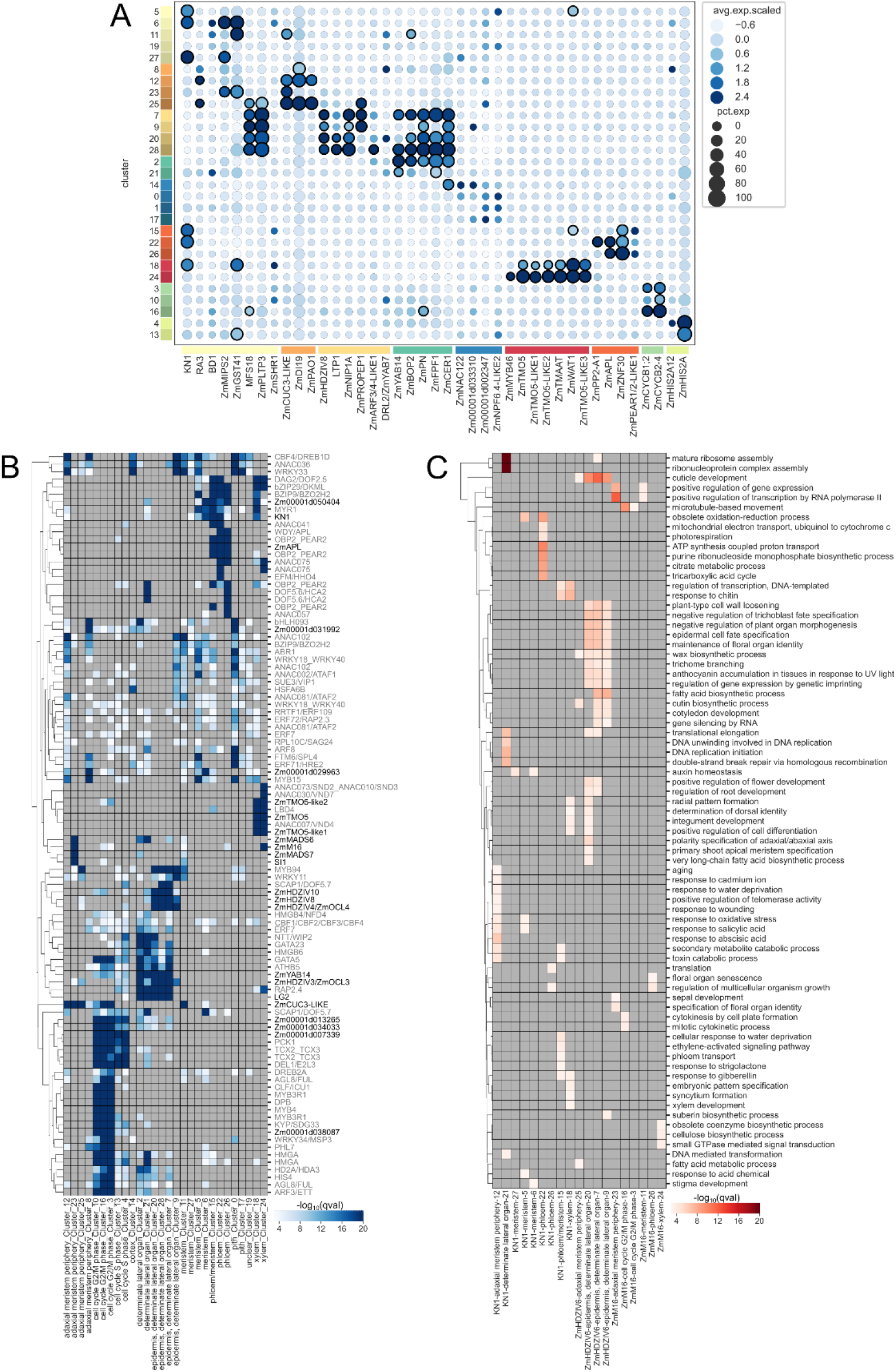
Assessment of the maize developing ear dataset and inferred regulons. (A) Bubble plot showing the expression of known marker genes from Xu et al., 2021 across the identified clusters. The color of the dots indicates the scaled average expression, while the size of the dots indicates the percentage of cells within the cluster expressing the gene. Dots with a thicker outline indicate genes which were identified as upregulated in the cell cluster. Marker genes and clusters are color-coded according to their annotation. (B) Clustermap showing cluster-specificity (the −log_10_(qval) of the single-cell cluster enrichment) of the top 150 inferred regulons. Gene annotation was retrieved from Arabidopsis orthologs. When a maize gene had multiple Arabidopsis orthologs, all are reported. TFs colored in black depict known cell-type specific maize regulators. (C) Clustermap showing the functional enrichment (-log_10_(qval) of GO-enrichment) of the TGs for KN1, ZmHDZIV6 and ZmM16 regulons.

**Supplemental Table 1. GWAS representation within regulons TGs.** The total upregulated-GWAS is 187. The percentages of column F are calculated as column E/column D, while the percentages of column G are calculated as column E/187.

**Supplemental Table 2. Overview of TF ChIP-Seq datasets used for the identification of optimal numbers of matches per tool and motif.**

**Supplemental Table 3. Overview of the datasets used to construct the Interactions-Gold Standard.**

**Supplemental Dataset 1. Overview of inferred regulons for the Denyer et al., 2019 Arabidopsis root dataset.** Regulons are ranked according to the global weighted Borda ranking obtained combining cluster-specificity, functional-specificity, closeness and betweenness centrality. The table reports A) TF geneID, B) alias, C) whether the TF is associated to a GO term of interest (relevant_known_TF), to any other GO term (known_TF), or to none (unknown_TF), D) GO terms associated to the TF, E) description of the GO terms, F) cell cluster the regulon acts, G) the corresponding cell-type, H) whether the TF is upregulated in the cell cluster (1) or just expressed (0), I) total number of regulons in the cell cluster, J) number of TGs controlled by the regulon, K) FDR-corrected p-value of cell cluster enrichment (cluster-specificity), L) out-degree centrality, M) closeness, N) betweenness, O) FDR-corrected p-value of functional enrichment of the regulon’s TGs (functional specificity - considering the lowest p-value for relevant terms), P) relevant GO term for which the regulon’s TGs show the most significant enrichment, Q) description of the GO term, R) number of TGs associated to the GO term, S) global Borda rank (over all regulons), T) local Borda rank (over all regulon of the cell-cluster).

**Supplemental Dataset 2. Overview of inferred regulons for the Wendrich et al., 2020 Arabidopsis root dataset.** Regulons are ranked according to the global weighted Borda ranking obtained combining cluster-specificity, functional-specificity, closeness and betweenness centrality. The table reports A) TF geneID, B) alias, C) whether the TF is associated to a GO term of interest (relevant_known_TF), to any other GO term (known_TF), or to none (unknown_TF), D) GO terms associated to the TF, E) description of the GO terms, F) cell cluster the regulon acts, G) the corresponding cell-type, H) whether the TF is upregulated in the cell cluster (1) or just expressed (0), I) total number of regulons in the cell cluster, J) number of TGs controlled by the regulon, K) FDR-corrected p-value of cell cluster enrichment (cluster-specificity), L) out-degree centrality, M) closeness, N) betweenness, O) FDR-corrected p-value of functional enrichment of the regulon’s TGs (functional specificity - considering the lowest p-value for relevant terms), P) relevant GO term for which the regulon’s TGs show the most significant enrichment, Q) description of the GO term, R) number of TGs associated to the GO term, S) global Borda rank (over all regulons), T) local Borda rank (over all regulon of the cell-cluster).

**Supplemental Dataset 3. Overview of inferred regulons for the Kim et al., 2021 Arabidopsis leaf dataset.** Regulons are ranked according to the global weighted Borda ranking obtained combining cluster-specificity, functional-specificity, closeness and betweenness centrality. The table reports A) TF geneID, B) alias, C) whether the TF is associated to a GO term of interest (relevant_known_TF), to any other GO term (known_TF), or to none (unknown_TF), D) GO terms associated to the TF, E) description of the GO terms, F) cell cluster the regulon acts, G) the corresponding cell-type, H) whether the TF is upregulated in the cell cluster (1) or just expressed (0), I) total number of regulons in the cell cluster, J) number of TGs controlled by the regulon, K) FDR-corrected p-value of cell cluster enrichment (cluster-specificity), L) out-degree centrality, M) closeness, N) betweenness, O) FDR-corrected p-value of functional enrichment of the regulon’s TGs (functional specificity - considering the lowest p-value for relevant terms), P) relevant GO term for which the regulon’s TGs show the most significant enrichment, Q) description of the GO term, R) number of TGs associated to the GO term, S) global Borda rank (over all regulons), T) local Borda rank (over all regulon of the cell-cluster), U) reference for functional annotation (PMID).

**Supplemental Dataset 4. Overview of inferred regulons for the rice root dataset.** Regulons are ranked according to the global weighted Borda ranking obtained combining cluster-specificity, functional-specificity, out-degree, closeness and betweenness centrality. The table reports A) TF geneID, B) alias, C) whether the TF is associated to a GO term of interest (relevant_known_TF), to any other GO term (known_TF), or to none (unknown_TF), D) GO terms associated to the TF, E) description of the GO terms, F) cell cluster the regulon acts, G) the corresponding cell-type, H) whether the TF is upregulated in the cell cluster (1) or just expressed (0), I) total number of regulons in the cell cluster, J) number of TGs controlled by the regulon, K) FDR-corrected p-value of cell cluster enrichment (cluster-specificity), L) out-degree centrality, M) closeness, N) betweenness, O) FDR-corrected p-value of functional enrichment of the regulon’s TGs (functional specificity - considering the lowest p-value for relevant terms), P) relevant GO term for which the regulon’s TGs show the most significant enrichment, Q) description of the GO term, R) number of TGs associated to the GO term, S) global Borda rank (over all regulons), T) local Borda rank (over all regulon of the cell-cluster).

**Supplemental Dataset 5. Overview of inferred regulons for the maize developing ear dataset.** Regulons are ranked according to the global weighted Borda ranking obtained combining cluster-specificity, functional-specificity, out-degree, closeness and betweenness centrality. The table reports A) TF geneID, B) alias, C) whether the TF is associated to a GO term of interest (relevant_known_TF), to any other GO term (known_TF), or to none (unknown_TF), D) GO terms associated to the TF, E) description of the GO terms, F) cell cluster the regulon acts, G) the corresponding cell-type, H) whether the TF is upregulated in the cell cluster (1) or just expressed (0), I) total number of regulons in the cell cluster, J) number of TGs controlled by the regulon, K) FDR-corrected p-value of cell cluster enrichment (cluster-specificity), L) out-degree centrality, M) closeness, N) betweenness, O) FDR-corrected p-value of functional enrichment of the regulon’s TGs (functional specificity - considering the lowest p-value for relevant terms), P) relevant GO term for which the regulon’s TGs show the most significant enrichment, Q) description of the GO term, R) number of TGs associated to the GO term, S) global Borda rank (over all regulons), T) local Borda rank (over all regulon of the cell-cluster).

